# Artemisinin-resistant *Plasmodium falciparum* Kelch13 mutant proteins display reduced heme-binding affinity and decreased artemisinin activation

**DOI:** 10.1101/2024.01.23.576340

**Authors:** Abdur Rahman, Sabahat Tamseel, Romain Coppée, Smritikana Dutta, Nawaal Khan, Mohammad Faaiz, Harshita Rastogi, Jyoti Rani Nath, Pramit Chowdhury, Ashish, Jérôme Clain, Souvik Bhattacharjee

## Abstract

The rapid emergence of artemisinin resistance (ART-R) poses a challenge to global malaria control efforts. ART potency is triggered by ferrous iron- and/or heme-mediated cleavage of the endoperoxide bond to generate reactive heme-ART alkoxy radicals and covalent heme-ART adducts that alkylate parasite targets or inhibit the detoxification of heme into β-hematin crystals; both of which lead to parasite death. Mutations in the *P. falciparum* Kelch-containing protein Kelch13 (PfKekch13) confer clinical ART-R, in which the resistant parasites exhibit impaired hemoglobin uptake, reduced heme yield, and thus decreased ART activation. However, a more direct involvement of PfKelch13 in heme-mediated ART activation has not been reported. Here, we show that recombinant, purified PfKelch13 wild-type (WT) protein displays measurable binding affinity for both iron and heme, the main effectors for ART activation. Comparative biochemical analyses further indicate weaker heme-binding affinities in the two Southeast Asian ART-R PfKelch13 mutants C580Y and R539T compared to the ART-sensitive WT and A578S mutant proteins, which ultimately translates into reduced yield of heme-ART derivatives. In conclusion, this study provides the first evidence for regulated ART activation *via* the heme-binding propensity of PfKelch13, which may contribute towards modulating the level of ART-R in malaria parasites with PfKelch13 mutations.

## Introduction

Malaria remains a major global health problem, with 249 million cases and 608,000 deaths reported in 2022 in the 84 malaria-endemic countries^1^. Artemisinin (ART) derivatives are key components of ART-based combination therapies (ACT), the current frontline treatment for uncomplicated malaria. ART derivatives contain an endoperoxide bridge that is cleaved within the malaria parasite by newly released hemoglobin-derived iron sources (predominantly ferrous iron, ferrous protoporphyrin IX (FPIX)/heme, or both) to form activated oxy-centered and C4-centered radicals^2^. These target the C-C, C-H, and other bonds in parasite proteins/lipids, and the resulting alkylation leads to extensive molecular damage and parasite death^3–7,8^. The alkylation of protein targets by ART radicals appears to be rather random, as two different click-chemistry studies using ART probes found little overlap in the set of alkylated parasite proteins^6,9^. Other anti-parasitic mechanisms include inhibition of β-hematin crystallization and heme detoxification by heme-drug adducts and, paradoxically, of endocytosis-mediated uptake of hemoglobin, the source of ART activator^10,11^. The role of heme-ART adducts is further supported by the detection of those adducts by LC-MS in live malaria parasites grown under culture conditions^12,13^. Non-heme (inorganic) iron, heme and hemoglobin are all implicated as key molecules in ART activation^14^, and selective inhibition of the heme degradation pathway or increasing heme levels (by exogenous addition of heme precursors) has been reported to reduce parasite sensitivity to ART or antagonize ART action, respectively^15,16^. In contrast, iron chelators antagonized ART efficacy, although they likely affected the iron molecule in FPIX/heme^7,14,17^. Recent observations using the click chemistry-based alkyne-labeled ART derivative and the free iron chelator deferoxamine have largely ruled out Fe^2+^ as an activator of ART^6,7,14,17^.

Unfortunately, the emergence and spread of resistance to ARTs in *Plasmodium falciparum* threatens the concerted progress made in reducing the global malaria burden over the past two decades following the introduction of ACTs. First reported in the Greater Mekong Subregion in 2008^18,19^, *P. falciparum* parasites with reduced susceptibility to ARTs are now widely reported in several regions of the world^20^. Resistance to ART is defined either clinically by a delayed parasite clearance half-life *in vivo* or by increased parasite survival *in vitro* after a brief exposure of early ring-stage parasites (0-3 h post erythrocyte invasion) to a high dose of dihydroartemisinin (DHA) in the ring-stage survival assay (RSA)^19,21^. Currently, mutations in the *P. falciparum* Kelch13 (PfKelch13) protein are the only confirmed determinants of clinical ART resistance^22,23^. PfKelch13 is a 726 amino acid protein encoded by the *pf3d7_1343700* gene on chromosome 13 and contains three structurally conserved domains: a coiled-coil-containing (CCC) domain, a broad complex, tramtrack and bric-à-brac (BTB) / poxvirus and zinc finger (POZ) domain, and a kelch domain (also known as Kelch-repeat propeller; KRP) which encompasses almost all the clinical ART resistance mutations^22,24^. The PfKelch13 C580Y and R539T mutants are the most common ART-resistant strains in Southeast Asia^25,26^, while the ART-R C469Y, R561H, R622I, and A675V mutants have emerged more recently in East Africa^27–29^. The A578S mutation is also common throughout Africa but does not confer ART resistance^25,26^.

Several studies have shown reduced steady-state cellular levels of PfKelch13 proteins in ART-resistant C580Y and R539T mutant parasites^30,31,32,33^. Then, conditional mislocalization of PfKelch13 using the diCre-based system resulted in developmental arrest of the replicating intraerythrocytic parasite at the ring-stage, highlighting the essential role of PfKelch13 at a stage where the ART-resistant phenotype also manifests^21,34,35^. GFP-tagged and native PfKelch13 were found to localize to membranous compartments proximal to the parasite food vacuole (FV) and to cytostomal structures near the parasite periphery^30,31^. Both mislocalized and ART-resistant PfKelch13 mutants exhibit reduced endocytic uptake of host cell material and free heme level at the ring stage^30,32^. It is now recognized that PfKelch13, together with other proteins, regulates hemoglobin endocytosis and subsequent degradation, thereby modulating the level of ART activation and toxicity. PfKelch13 also colocalizes with parasite endoplasmic reticulum, Rab-positive vesicles, and partially with parasite mitochondria upon DHA treatment^36^. Of note, three PfKelch13-associated malarial structures (cytostome, FV, and mitochondria) are liked to either heme-loaded hemoglobin capture and degradation or heme synthesis. Cytostome-derived vesicles loaded with host hemoglobin may also contain newly released free heme from early vesicular degradation of hemoglobin^37^. The parasite mitochondrion, together with the plastid, are sites of *de novo* heme synthesis, and mitochondrial metabolism can modulate ART resistance. Indeed, deletion of the malaria mitochondrial protease PfDegP has been reported to reduce heme levels and DHA susceptibility^38^. Finally, large amounts of heme are released during the digestion of hemoglobin in the food vacuole^39,40,41^. Thus, the detection of PfKelch13 proximal to heme/hemoglobin-containing cellular structures (either cytosolic or membrane-bound) and the evidence of a labile heme pool (∼1.6 μM) that is stably maintained in the parasite^42^ raises the possibility of direct interaction(s) between PfKelch13 and heme/hemoglobin. Since both hemoglobin uptake and heme are involved in ART activation, are targets of activated ART, and are regulated by the ART-R determinant PfKelch13, we hypothesize a more direct link between heme and PfKelch13. In the literature, we identified one KRP domain with iron binding properties from the field pennycress *Thlaspi arvense* thiocyanate-forming protein (TaTFP) of the *Brassicaceae* family^43,44^. Previous studies have shown increased activity of TaTFP upon Fe^2+^ supplementation compared to Fe^3+^, while the other metal ions were ineffective^43–46^. Here, we sought to test the hypothesis that PfKelch13 could interact with iron and heme. Recently, we reported the successful purification of recombinant PfKelch13 WT and A578S, C580Y, and R539T mutant proteins expressed in *Escherichia coli*^47^. These recombinants led us to investigate molecular interactions between PfKelch13 and putative activators of ART, namely iron and heme. We present the first biochemical evidence along with *in silico* insights for the differential heme-binding affinity between ART-sensitive (WT and A578S) PfKelch13 proteins and the two ART-resistant mutants (C580Y and R539T) and their effect on heme-ART adduct yields.

## Results

### The KRP domain of PfKelch13 shares structural features with the iron-binding KRP domain of

#### Thlaspi arvense TFP

First, we compared the KRP domain of PfKelch13 with that of the enzymatically active, iron-binding KRP of the field pennycress *Thlaspi arvense* thiocyanate-forming protein (TaTFP) of the *Brassicaceae* family^43,44^ (**Figure 1A**). The binding of TaTFP to iron directly involves amino acid positions of the KRP domain (E266, D270 and H274, while W309 supported the positioning of H274)^48,49^. By performing pairwise sequence and 3D structure alignments of the KRP domains of PfKelch13 and TaTFP, amino acid E266 of TaTFP aligned in both sequence and structure with E688 of PfKelch13, with a similar side chain orientation and located in the shallow pocket of KRP (**Figure 1B-D** and **Supplementary Figure S1**). Among the other amino acids, W706 of PfKelch13 only showed alignment in sequence with W309 of TaTFP; while the other amino acids, *i.e.,* E691 and H697 of PfKelch13, did not align in either sequence or structure with D270 and H274 of TaTFP (**Figure 1B-D**). We also searched *in silico* for putative iron (Fe^2+^ or Fe^3+^)-interacting residues in the amino acid sequences from the full length PfKelch13, the truncated PfKelch13 (TrK13-WT; as reported in our previous study^47^ with the amino acids coding for the step-tag sequence at the C-terminus) using the MIB server (http://bioinfo.cmu.edu.tw/MIB/)^50^. Typically, 95 and 117 amino acid residues in the full length PfKelch13, and 97 and 110 different amino acid residues in TrK13-WT were predicted to bind Fe^2+^ and Fe^3+^, respectively, including six and three residues from the strep affinity tag in TrK13-WT (summarized in **Supplementary Table S1**). Taken together, these results prompted us to biochemically test the binding ability of different metal ions to recombinant PfKelch13.

**Figure 1.**
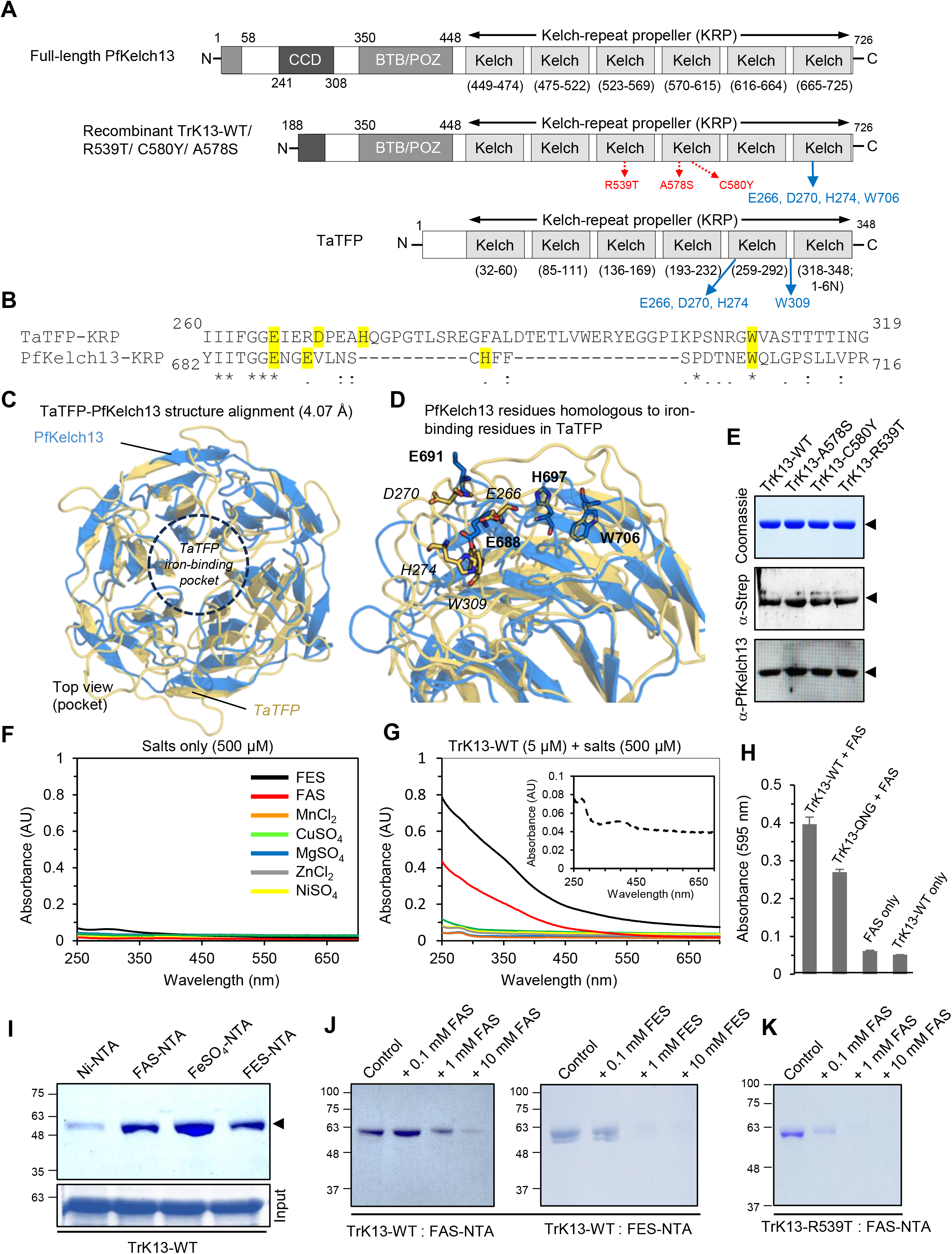
Structural and functional similarities in the iron-binding properties between the KRP domains of PfKelch13 and TaTFP. **A.** Schematic representation of the domain organization in full-length PfKelch13 (top), recombinant TrK13-WT (middle), and TaTFP (bottom), showing the respective spans of the coiled-coil domain (CCD), bric-à-brac; (BTB)/poxvirus and zinc-finger (POZ) domain (BTB/POZ), and the Kelch-repeat propeller (KRP) domain. The three specific mutations in the full-length PfKelch13: R539T, C580Y and A578S are indicated in red text. The iron binding residues in TaTFP and corresponding homologous amino acids in TrK13 are shown in blue text. **B.** Sequence alignment between a part of the KRP domains of TaTFP and PfKelch13. Residues highlighted in yellow represent positional conservation between PfKelch13 and those in TaTFP involved in Fe^2+^ binding. **C-D.** Structural alignment of the PfKelch13 propeller (blue) and TaTFP (gold) proteins. Protein structures are shown as cartoons. Iron-binding amino acid residues in TaTFP (italic) and homologous sites in PfKelch13 (bold) are shown as sticks (in D). **E.** SDS-PAGE and Western blot profiles showing affinity-purified TrK13 proteins of ∼61 kDa (arrowheads) and their specific recognition by antibodies. **F-G.** UV-visible spectroscopy scans showing the increase in absorbance profiles of TrK13-WT only in the presence of FAS and FES, but not with other metal salts (G). The corresponding spectrum of free salts in the absence of any TrK13-WT protein is shown in F. The spectrum profile of TrK13-WT protein without any metal salt is shown as an inset. The graphs shown are best representative of three independent experiments. **H.** Graph showing the relatively higher binding of the TrK13-WT protein to FAS, as measured by absorbance at 595 nm using the Ferene S assay, compared to the TrK13-QNG mutant protein. Data with their respective standard errors from three independent experiments are shown. **I.** SDS-PAGE showing higher affinity of TrK13-WT protein (arrowhead) for FAS-, FeSO4- and FES-NTA beads compared to Ni-NTA beads. Input samples are shown below. **J-K.** Inhibition of TrK13-WT (J) or TrK13-R539T (K) binding to FAS-NTA (J; left) or FES-NTA (J; right) beads with stepwise increase of free FAS (J; left gel) or FES (J; right gel) concentration. Molecular weight standards (in kDa) for all SDS-PAGE gels are shown on the left.

### The recombinant PfKelch13 protein shows specific binding to iron salts but not to salts of other divalent cations

The cloning, expression and purification of recombinant PfKelch13 proteins from *E. coli* were performed as described in our previous study^47^. Similar to our earlier observations, the recombinant purified PfKelch13 was deleted in the Apicomplexa-specific N-terminal segment (TrK13) and all the TrK13 variants (WT and R539T, A578S and C580Y mutants) migrated as a ∼61 kDa band in SDS-PAGE (**Figure 1E** and **Supplementary Figure S2**). TrK13 proteins were immunodetected in western blots using either commercially available antibodies against the strep tag or custom-generated antibodies against a PfKelch13 peptide^47^. We then looked for evidence of iron (and then heme) binding using different methods. First, TrK13 proteins (5 μM) incubated in solution with 100-fold molar excess (500 μM) of individual metal salts (MnCl_2_, CuSO_4_, MgSO_4_, ZnCl_2_, NiSO_4_, ferrous ammonium sulfate or FAS, and ferric ammonium sulfate or FES) were analyzed by UV-visible spectrophotometry. The free metal salts alone showed no observable absorbance in the 250-700 nm spectrum at these concentrations (**Figure 1F**). Interestingly, TrK13-WT showed a dramatic increase in absorbance at all wavelengths in the presence of the two iron salts FAS and FES (**Figure 1G**). At 280 nm (representing the cumulative absorbance by the aromatic amino acids tyrosines, tryptophans, and phenylalanines), we observed a 5.8-fold and 12.8-fold increase in the absorbance of TrK13-WT in the presence of FAS and FES, respectively, compared to TrK13-WT alone (no salts). The solubility of TrK13-WT remained unchanged in the absence or presence of FAS or FES at these concentrations (**Supplementary Figure S3**). This FAS- and FES-dependent increase in absorbance was a proxy for ligand binding and was also detected in TrK13-C580Y, TrK13-R539T and TrK13-A578S mutant proteins (**Supplementary Figure S4**). These data suggest that TrK13 binds iron salts and that the three mutations tested do not significantly alter the binding.

Since the major source of iron in the mature erythrocytes is the heme in hemoglobin, which contains iron in Fe^2+^ redox state^51^, we investigated the binding of TrK13-WT to FAS using the *in-solution* Ferene S colorimetric assay, in which iron is reduced to Fe^2+^ by the addition of ascorbic acid, followed by bidentate chelation of Fe^2+^ by Ferene to form a stable blue complex between pH 3.0 and 6.0^52^. The TrK13-WT protein (5 μM) was incubated with a 30-fold molar excess (150 μM) of FAS, and the mixture was then desalted by size-exclusion chromatography using PD Spintrap™ G-25 desalting column. An approximately 10-fold increase in the absorbance at 595 nm was measured in the eluate for TrK13-WT incubated with FAS, as compared to the FAS only or TrK13-WT only controls that were similarly processed (**Figure 1H**). Based on the sequence alignment between TaTFP and PfKelch13, we also included a recombinant TrK13 variant (TrK13-QNG) with E688Q-E691N-H697G point mutations at the three positions homologous to the Fe^2+^ binding positions in TaTFP^48,49^. We observed ∼33.2% reduction in absorbance at 595 nm in the TrK13-QNG variant compared to TrK13-WT. Taken together, these results indicate that FAS molecules were co-purified with the TrK13 proteins, suggesting iron binding. Since the previous *in silico* analysis predicted a hundred of Fe^2+^-binding sites in TrK13-WT, we did not proceed to generate other mutants.

We further evaluated the binding of TrK13-WT to iron by using immobilized-metal affinity chromatography, in which the metal ion is coupled to a resin matrix *via* nitrilotriacetic acid (NTA). We tested three different immobilized iron matrices (*i.e.*, FAS-NTA, FeSO_4_-NTA, and FES-NTA; collectively referred to as Fe-NTA)^53,54^ and Ni-NTA as a control. TrK13-WT (5 µM) was incubated with the different matrices in neutral pH buffer under rotating conditions, and after extensive washing, the matrix-bound proteins were eluted with Laemmli buffer and separated by SDS-PAGE. After Coomassie staining, the ∼61 kDa band corresponding to TrK13-WT was ∼3 to 5 fold more intense with the three Fe-NTA matrices compared to the Ni-NTA control (**Figure 1I**). We also observed less eluted TrK13-WT when excess free FAS or FES (0.1 to 10 mM) was added during the bead incubation period, consistent with specific binding (**Figure 1J**). A similar trend was observed for the ART-R mutant TrK13-R539T (**Figure 1K**).

In our earlier study, we reported the homo-hexameric assembly of TrK13-WT by SAXS, which remained stable in the presence of up to 2 M urea concentration. At higher urea concentrations, the hexameric TrK13-WT assembly was found to dissociate into ∼61 kDa monomeric units^47^. We therefore tested the binding of TrK13-WT to FAS-NTA in the presence of increasing urea concentrations from 0 to 6 M. Compared to no urea, there was an approximately 52% reduction in the amount of eluted TrK13-WT at urea concentrations of 2 M and beyond (**Supplementary Figure S5**). Since the monomeric TrK13-WT unit still retained significant FAS-NTA-binding, this suggests that iron binding is not strictly dependent on the hexameric assembly of the TrK13-WT protein.

### The recombinant WT and mutant TrK13 proteins show specific interactions with heme

We then investigated whether TrK13 also binds to the iron-containing heme molecule. Whenever possible, we included the ART-R TrK13-R539T and TrK13-C580Y and the ART-S TrK13-A578S recombinant proteins.

First, we qualitatively evaluated the heme-binding potential of TrK13-WT by *in gel* staining^55,56,57^ using 3,3′,5,5′ tetramethylbenzidine (TMBZ). After incubation of 10 µM TrK13-WT with equimolar heme (hemin chloride reduced to heme in the presence of 2 mM sodium dithionite) for 30 min, the mixture was resolved by native PAGE. After gel staining with TMBZ, an intense bluish-green band was detected at >180 kDa (**Figure 2A**), a molecular weight reminiscent of the TrK13-WT oligomer reported previously^47^. Without heme co-incubation, no TMBZ staining was observed for either the TrK13-WT protein or the proteins from the molecular weight standards. The light bluish-green smear tail was likely due, at least in part, to background TMBZ staining^55^. The parallel Coomassie stained replicate gel showed a similar staining pattern (**Figure 2A**). These data indicate that heme molecules were co-migrating with the TrK13-WT protein.

**Figure 2.**
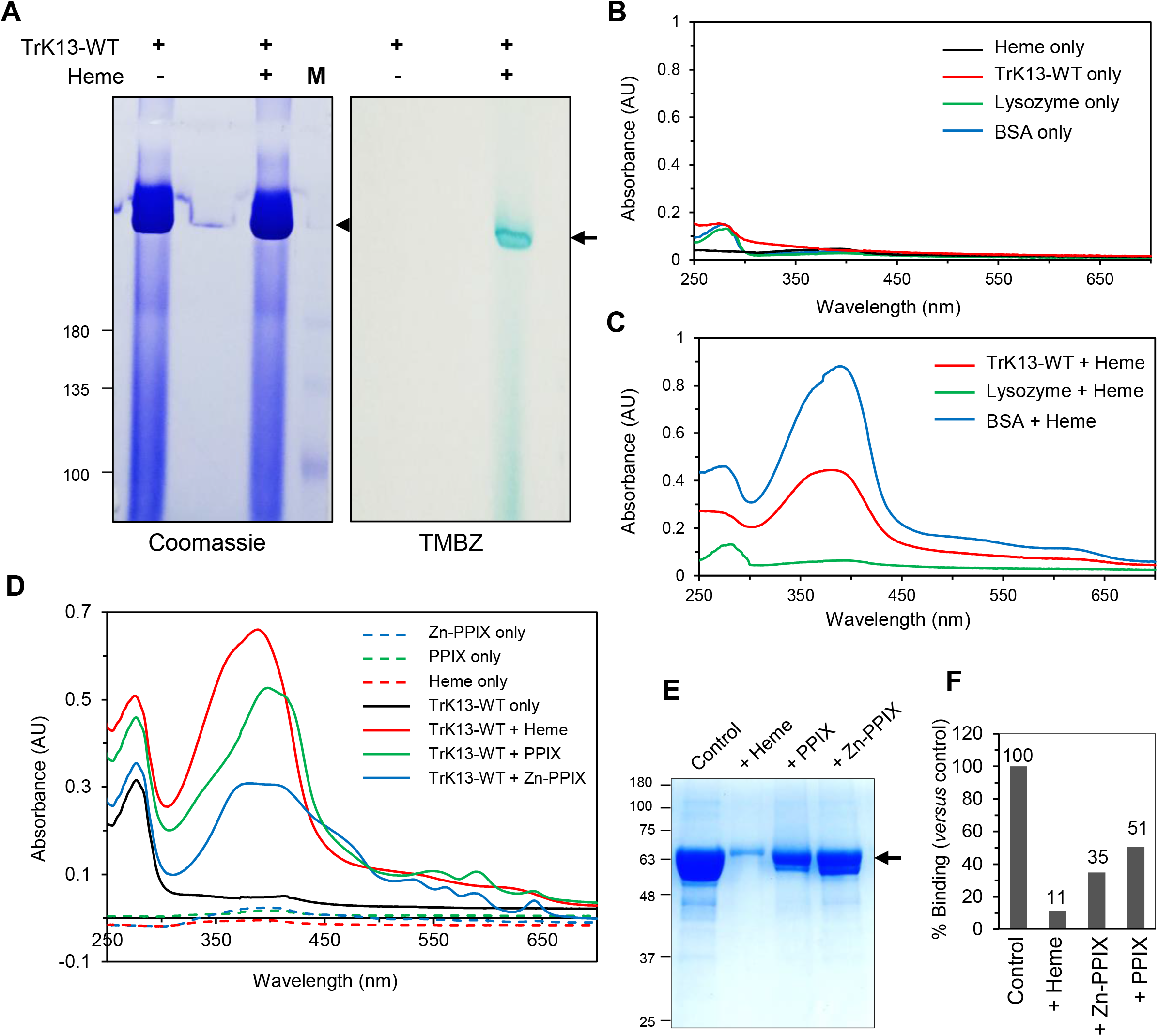
The recombinant TrK13-WT protein shows a specific interaction with heme/FPIX. **A.** Native PAGE of TrK13-WT shows a bluish-green TMBZ-positive band (arrow, right gel) only in the heme-preincubated sample (right lane) but not in the absence of heme (left lane). Corresponding Coomassie-stained SDS-PAGE of TrK13-WT protein with or without preincubated heme are shown on the left gel. **B-C.** UV-visible spectroscopy scans showing increase in the relative absorbance profiles of PD-10 desalted TrK13-WT (red) and positive control BSA (blue), but not negative control lysozyme (green), in the presence of heme (B) compared to their absorbance profiles without heme (C). No soret peak of free heme (black) is detected in C at this concentration (10 μM), indicating effective heme removal by the G25 desalting column. **D.** UV-visible spectroscopy scans showing the increase in the relative absorbance profiles of G-25 desalted TrK13-WT protein (10 μM) in the presence of 10 μM heme/FPIX (solid red line) or its analogs such as Zn-PPIX (solid blue line) or PPIX (solid green line) compared to the control (solid black line). The spectral profiles of the remaining unbound heme variants after the G-25 desalting step are shown as corresponding colored dotted lines and indicate efficient removal as no Soret peak was visible. **E-F.** SDS-PAGE (E) and graph (F) showing the relative abundance of TrK13-WT protein (arrow) pulled down with hemin-agarose beads in the absence or presence of excess free heme, PPIX, or Zn-PPIX. Molecular weight standards (in kDa) for all SDS-PAGE images are shown on the left.

We then evaluated the binding of heme to WT and three mutant TrK13 proteins by using immobilized heme affinity chromatography with hemin-agarose beads^58^. After extensive washing, the bead-bound proteins were eluted with Laemmli buffer and resolved by SDS-PAGE. All four TrK13 were detected in the Coomassie-stained gel (**Supplementary Figure S6**). The elution yield was ∼30-40% of the initial input for each TrK13 variant, suggesting no detectable major effect of the mutations on the hemin-agarose binding property of the TrK13 proteins. Note that the hemin-agarose experiments were limited by the fixed heme concentration in the beads, as well as the possibility of steric constraints on heme accessibility by the proteins.

To overcome these limitations, the TrK13-WT protein (10 µM) was incubated with an equimolar concentration of heme (reduced with sodium dithionite) for 10 min and the mixture was desalted using the G-25 columns. The UV-visible spectral profile of the desalted mixture showed a large increase in absorbance throughout the 250-700 nm wavelength spectrum with a prominent Soret peak, typical for heme-containing molecules (**Figure 2C**), compared to the desalted TrK13-WT alone control (**Figure 2B**). Our desalting procedure using G-25 column efficiently removed the free heme, as no Soret peak was visible after desalting the heme alone sample (**Figure 2B***; the spectrum scan of free heme without desalting is shown in Supplementary figure S9A*). The TrK13-WT-heme mixture showed a hypsochromic or blue shift from 410 nm to 370 nm, corresponding to the Soret peak. Of the two control proteins incubated with heme and processed similarly, BSA (a confirmed heme-binding protein^59–61^) showed a larger Soret peak, whereas lysozyme (which has no heme-binding affinity^62^) showed no increase in absorbance (**Figures 2C**).

### The TrK13-WT-heme interaction involves both the Fe^2+^ ion and the PPIX porphine core

The heme molecule consists of protoporphyrin IX (PPIX), a tetrapyrrole macrocyclic porphine core with a Fe^2+^ cation in the center. To investigate whether the TrK13-WT heme interaction involves one or both components, TrK13-WT (10 μM) was incubated with equimolar concentrations of heme or heme analogs such as PPIX (without the Fe^2+^ ion) or Zn-PPIX (with Zn^2+^ replacing the Fe^2+^ cation), desalted through G-25 columns, and analyzed by UV-visible spectrophotometry. As previously observed for heme, there was an increase in the absorbance spectrum of TrK13-WT regardless of the heme analog tested. Specifically, distinct peaks with absorbance maxima between 300-450 nm wavelengths were visible in all cases (and absent in the protein-only control) and were characteristic of the heme analog used (**Figure 2D**). Efficient desalting was also confirmed by the absence of such maxima peaks in the control samples, thus excluding any contributions from free heme or its analogs. To confirm the binding of heme analogs, TrK13-WT (10 µM) was incubated with hemin-agarose beads in the presence of a 10-fold molar excess of heme analogs. After extensive washing, co-incubation with heme or analogs decreased the amount of TrK13-WT eluted (**Figures 2E-F**), with heme displaying the largest inhibition (∼90%) compared to PPIX (49%) and Zn-PPIX (65%). Taken together, these results strongly suggest the involvement of both the Fe^2+^ metal ion and the PPIX moiety of heme in TrK13-WT interactions.

### Kinetics of TrK13 tryptophan fluorescence quenching by heme

In tryptophan fluorescence quenching, the tryptophan residues within a protein are selectively excited at a wavelength of 295 nm and the emission spectrum, which peaks at 300-350 nm, is measured in the absence or presence of a ligand^63,64^. The full-length PfKelch13 has seven tryptophan residues at positions 229, 490, 518, 565, 611, 660 and 706, while the recombinant TrK13 proteins (WT and mutants) have nine tryptophan residues due to two additional tryptophan residues contributed by the one-strep tag fusion at the C-terminus (**Supplementary Figure S7**). Since the tryptophan residues are distributed along the entire stretch of TrK13 sequence, we examined the interactions of heme-TrK13 by fluorescence quenching. The fluorescence spectra of WT, R539T, C580Y, and A578S TrK13 proteins exhibited peak maxima between 332-337, 338-341, 336-338, and 326-333 nm, respectively, and were gradually quenched by increasing concentrations of heme (up to 5 µM; **Figure 3A-D**). Blue shifts to lower wavelengths (i.e., 329 nm) upon heme binding were observed for all TrK13 proteins, which could be attributed to the formation of a heme-protein complex accompanied by increased hydrophobicity around the tryptophan sites^59,65^. Interestingly, 50% quenching of tryptophan fluorescence was achieved at heme concentrations between 2-2.4 μM for the ART-sensitive TrK13-WT and -A578S proteins, while it was achieved at 0.9-1 μM heme for the ART-resistant mutants TrK13-R539T and -C580Y. The Stern-Volmer plots showed an upward curvature for all TrK13 proteins, especially at higher heme concentrations, suggesting either combined fluorescence quenching or the existence of >1 heme-binding sites with different accessibility to the tryptophan residues in TrK13 proteins^60^ (**Figure 3E**). Values for the Stern-Volmer quenching constant K_SV_ (determined only for the linear part of the Stern-Volmer plot^66^) excluded the effect of collisional quenching in water. Similarly, the values for the quenching rate constants were in the order of 10^13^, which is generally considered to be well above the maximum k_q_ due to collisional quenching^67^. Interestingly, we observed marked differences in the Stern-Volmer constants (K_SV_) between the TrK13 variants (**Figure 3E**). The two mutants associated with ART-R in Southeast Asia (TrK13-R539T and -C580Y) showed higher K_SV_ values (1.18 and 1.76 μM^-1^, respectively) compared to those of the ART-sensitive proteins TrK13-WT and -A578S (0.60 and 0.66 μM^-1^, respectively). The quenching mechanism can be attributed to FRET arising from appreciable overlap between the tryptophan emission and heme absorption. TrK13-WT showed the least quenching while for the mutants, there was an overall increase in quenching with TrK13-C580Y showing the maximum. Since the extent of energy transfer is dependent on the distance between the donor (tryptophan residue) and the acceptor (heme), the quenching data suggest that heme binding brings about definite changes in the protein structure/conformation that lead to changes in the distance between Trp and heme. These results were in good agreement with our earlier structural studies of TrK13 proteins using SAXS, which indicated an increased length of the oligomeric assembly and a more skewed orientation of the KRP domains away from the long axis of the protein structures in TrK13-R539T and -C580Y mutants compared to TrK13-WT and -A578S mutant^47^. However, quantitative assessment of the true heme-binding affinities of these recombinants was further estimated by more sensitive assays.

**Figure 3.**
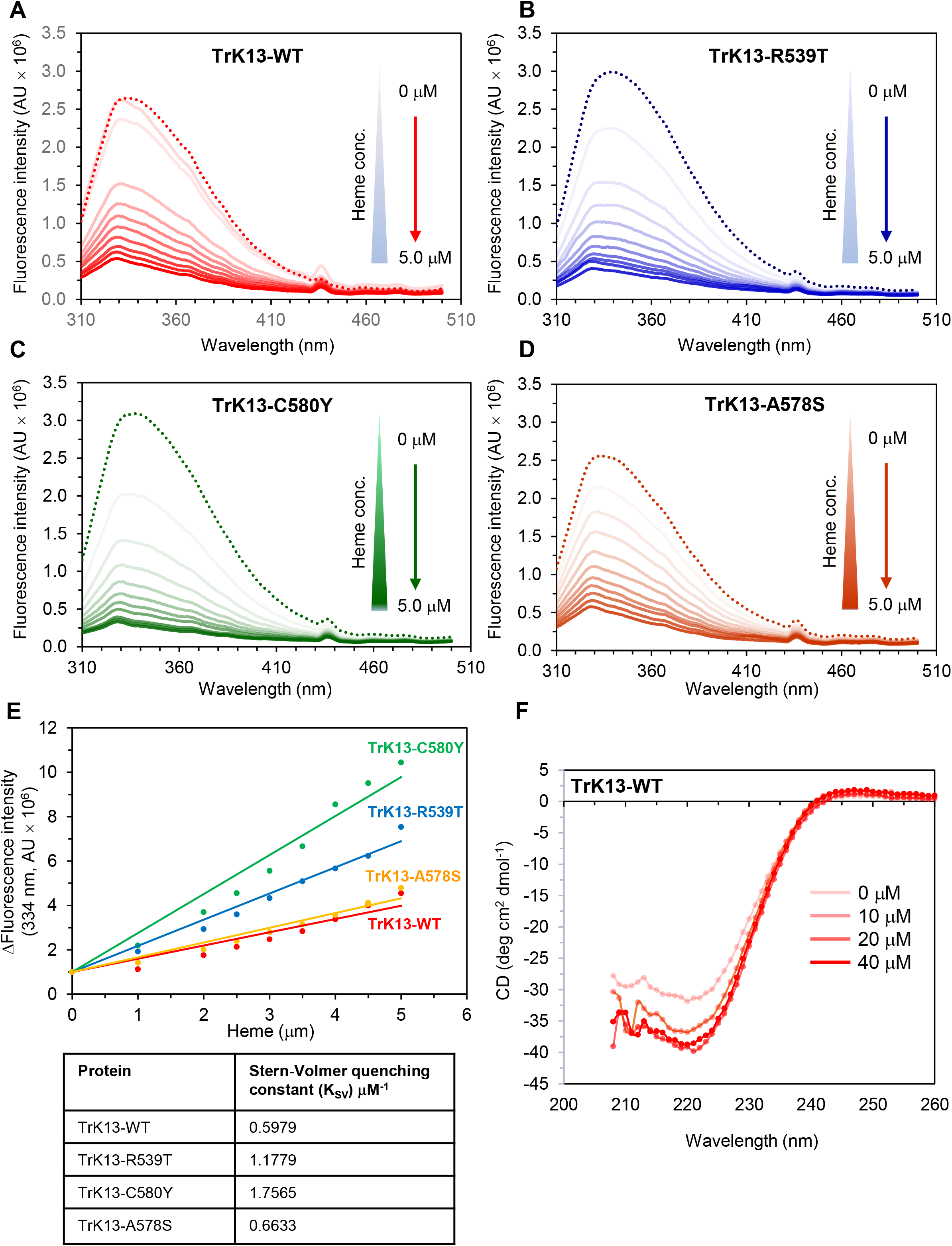
Heme-induced fluorescence quenching of tryptophan residues in the four variants of TrK13 proteins without profound changes in the circular dichroism profile. **A.-D.** Fluorescence spectra of TrK13-WT (A), TrK13-R539T (B), TrK13-C580Y (C), and TrK13-A578S (D) incubated in the absence (dotted lines) or presence of heme at final concentrations between 0.5-5.0 μM heme (gradual light to dark shading in each color) and showing a successive decrease in fluorescence intensity. **E.** Changes in tryptophan fluorescence quenching in the four variants of TrK13 as deduced from A-D. The Stern-Volmer quenching constant (K_SV_) was calculated from the equation F_0_/F = K_sv_ [Q] + 1, where F_0_ = fluorescence intensity in the absence of heme, F = fluorescence intensity in the presence of heme, [Q] = concentration of quencher (heme). Curves are the average of three replicates. **F.** Far-UV CD spectrum for titration of TrK13-WT protein with increasing concentrations of heme.

The effect of increasing concentrations of FPIX/heme on the secondary structure of TrK13-WT protein was measured by circular dichroism (CD) spectroscopy (**Figure 3F**). In the absence of heme, the CD profile revealed primarily a β-pleated secondary structure with traces of α-helical regions. In the presence of up to 20 μM heme, changes in secondary structure were observed with a gradual increase in molar ellipticity, particularly affecting the 222 nm minimum, suggesting an enhancement in helical content^68^. Not much change was observed at higher heme concentrations (40 μM). In summary, these studies indicated changes in the helical content, but not in the overall secondary structure of TrK13-WT upon heme-binding. The CD spectral changes upon ligand binding does not rule out heme binding, as binding sites can be located far from the aromatic amino side chains in the protein without causing a detectable change in secondary structure.

### Quantitative evaluation of TrK13-heme interactions using microscale thermophoresis

We quantitatively investigated the true affinity between heme and TrK13 using label-free microscale thermophoresis (MST), which exploits the intrinsic fluorescence of tryptophan residues of proteins^69^. When titrated against increasing concentrations of heme, the thermophoretic mobility of TrK13 changed, indicating effective interactions. The dose-response curves showed a negative response amplitude for all TrK13 variants, indicating a lower MST signal of the heme-protein complex compared to TrK13 alone (**Figure 4A-D**). Furthermore, equilibrium dissociation constant (K_d_) calculations revealed values of 61.5 ± 12.4 nM and 66.0 ± 17.9 nM for the ART-sensitive TrK13-WT and TrK13-A578S mutant respectively (**Figures 4E-F**). Interestingly, higher K_d_ values were calculated for the TrK13-R539T and TrK13-C580Y mutants (131.0 ± 14.0 nM and 204.0 ± 20.4 nM, respectively, **Figure 4F**). These results indicate a comparatively weaker association of heme with the two ART-R mutants compared to the ART-sensitive proteins.

**Figure 4.**
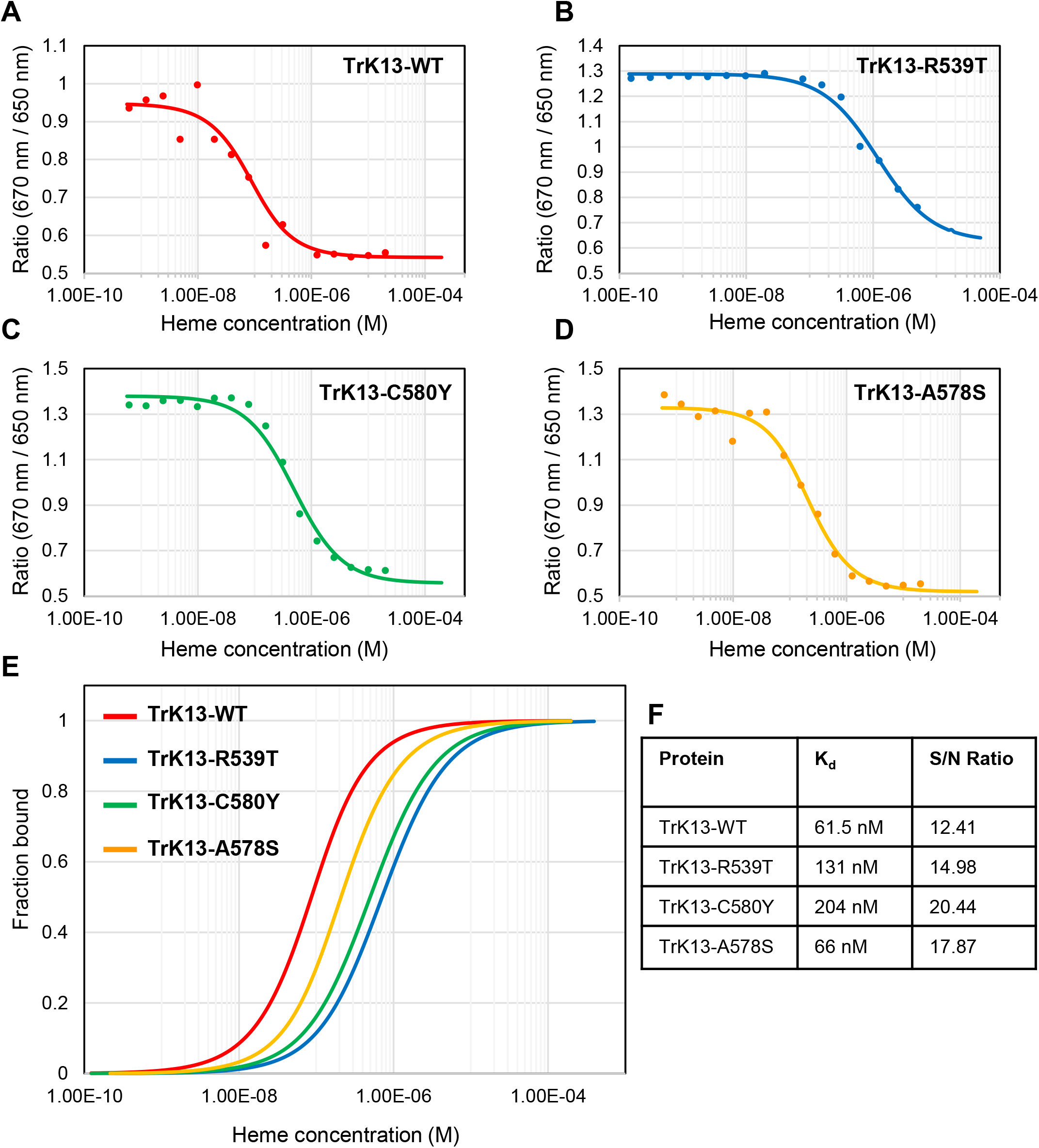
Quantitative evaluation of TrK13-heme binding by MST. **A.-D.** Dose-response curves showing negative amplitude of response for TrK13-WT (B), TrK13-R539T (C), TrK13-C580Y (D) and TrK13-A578S (E) titrated against increasing concentrations of heme. MST experiments were performed at medium MST power at 25°C. LED power was set to 60% (B and E) or 40% (C and D) excitation. **E.** Comparative dose-response curves for the binding interaction between the four TrK13 variants and heme, as indicated. The protein concentration was kept constant at either 50 nM (TrK13-WT or TrK13-A578S) or 25 nM (TrK13-R539T or TrK13-C580Y), while the ligand heme concentration varied from 0-5 µM. **F.** Table showing the decreasing affinity scale: TrK13-WT > TrK13-A578S > TrK13-R539T >TrK13-C580Y and signal-to-noise ratio.

### *In silico* docking and computational approaches predict heme interaction sites in PfKelch13

Next, we searched for putative PfKelch13-heme interaction sites using an *in silico* approach and the TrK13 3D structure^47^. No predicted cavity overlapped two monomers of the homo-hexameric assembly of TrK13-WT. Since disruption of the oligomerization state by high concentrations of urea abolished ∼50% of Fe-NTA binding (**Supplementary Figure S5**) or hemin-agarose binding (data not shown), we used the TrK13-WT monomeric unit to predict cavities, which were then individually docked to a heme molecule. Eleven heme-binding cavities were predicted on TrK13-WT monomeric unit including seven in the KRP domain (**Figure 5**). The binding cavities found in BTB and KRP were also predicted using the 1.81 Å resolution X-ray structure of WT PfKelch13 BTB-KRP (PDB ID: 4YY8)^70^, showing robustness to the source of the WT template structure. The top three predicted cavities with the lowest binding free energies with heme were located in the CCC (-7.9 kcal/mol), BTB (-7.6 kcal/mol), and KRP (-7.9 kcal/mol) domains. The highest affinity interaction in the KRP domain involved amino acid positions R529, R597, and E688, all located in the highly conserved shallow pocket^24^. Of note, the amino acid homologous to PfKelch13 E688 in the KRP domain of TaTFP (E266) is directly involved in iron-binding^48,49,71^. Two predicted TrK13-WT-heme interactions involved the amino acid positions R539 or C580, while the amino acid position A578 was never associated with any interaction (**Supplementary table S2**).

**Figure 5.**
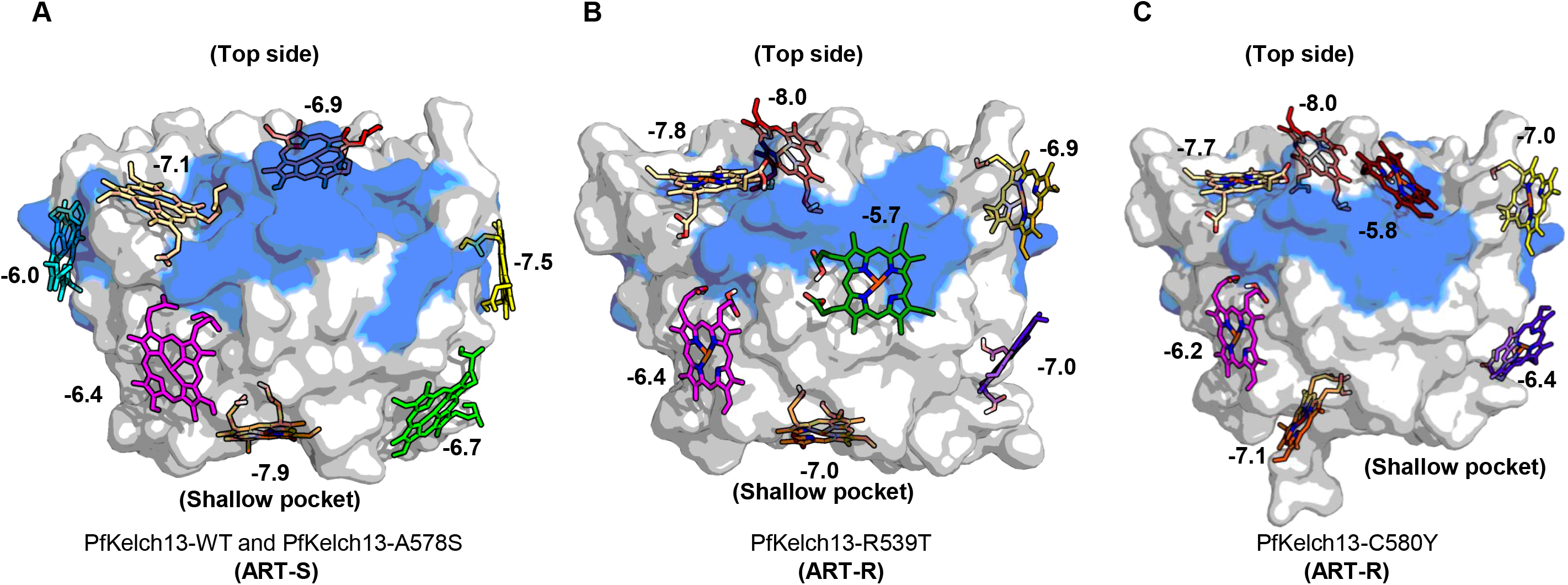
*In silico* molecular docking of heme to ART-S and ART-R variants in the KRP domain of PfKelch13. **A.-C.** Structures showing the top seven docked poses of heme in the KRP domains of (A) PfKelch13-WT and -A578S, (B) -C580Y, and (C) -R539T structures. The PfKelch13 structures are shown as surface and colored according to the secondary structure (beta-strand, blue; loop, gray). Heme is shown as stick. The binding affinity (expressed in kcal/mol) is given for each of the predicted heme-binding poses (the more negative the value, the better the binding affinity). Each pose is represented by a different color.

We then similarly tested *in silico* the three TrK13 mutant structures. For the TrK13-A578S mutant, whose structure was very similar to TrK13-WT using SAXS-derived data^47^, same cavities and heme binding affinities were predicted (**Figure 5**). In contrast, different predictions were found for TrK13-R539T and TrK13-C580Y, which have slightly different 3D structures compared to WT^47^. While several cavities of the mutants were common to the WT ones, we also detected new ones that were not found in the WT structure and *vice versa*. In these two mutant structures, the highest affinity interaction was located at the top of KRP (on the opposite side of the shallow pocket) with binding free energies of -8.0 kcal/mol. The predicted interactions remained in the shallow pocket, but with slightly lower affinities (TrK13-R539T: -7.0 kcal/mol; TrK13-C580Y: -7.1 kcal/mol; **Figure 5**) compared to the WT and A578S structures (-7.9 kcal/mol). These predicted interactions in the mutants again involved the amino acid position E688 (**Supplementary Table S2**). The mutated residue T539 in TrK13-R539T was directly involved in one of the predicted interactions (with a moderate score of -5.7 kcal/mol), while the mutated residue Y580 in TrK13-C580Y was no longer involved in any heme interaction. Overall, some regions of the KRP domain were predicted to interact with heme and some differences were observed between ART-R and ART-S proteins, consistent with these two groups of proteins exhibiting different binding affinities as measured experimentally.

### The TrK13-bound heme forms heme-ART activated species (AAS)

A general scheme for endoperoxide bond cleavage in ART derivatives by reduced heme to generate either oxy-radicals or C4-centered ART adducts has been reported previously^3,13,72,73^. Because we were unable to distinguish between the peaks corresponding to either oxy-radicals or C4-centered ART adducts in the downstream assays, we chose to use the term ‘ART activated species (AAS)’ to jointly represent the products. We investigated the formation of AAS under reducing conditions by high-performance liquid chromatography (HPLC) after 30 min incubation of heme with ART or analogs (**Supplementary Figures S8-S9**). The time-dependent formation of AAS in our experimental setup is shown in **Supplementary Figure S9B**. Heme-containing molecules were detected based on the presence of porphyrins excited at 390–410 nm (Soret peak) whereas ART and DHA have been previously reported to absorb only in the UV region between 190-210 nm^74^. At a molar ratio of 1:1.2 (heme: ART or respective derivatives), additional peaks with 5 to 10 times lower intensity and slightly delayed elution times were visible in the chromatogram at 410 nm with either ART or DHA compared to heme alone or heme with an inactive analog of ART such as deoxyartemisinin (DOA) (**Figure 6A**). The elution time and profile of these additional delayed peaks varied with the derivatives tested, namely DHA or ART (**Figure 6A**) or artesunate (AS, **Supplementary Figure S8**), and likely corresponded to AAS as reported in previous studies^12,13,10^. The formation of AAS was also verified by LC-MS in the *in vitro* reaction mixtures of heme with or without ART. As shown in other studies, prominent peaks at m/z 529.4, 543.4, 557.2, and 616.2 characteristic of heme and common degradation products generated by in-source fragmentation of heme and oxidation-reduction processes during electrospray ionization were observed in the heme-only sample (**Figure 6B**)^12,75^. When heme (under reducing conditions) was reacted *in vitro* with ART (mass of 282.3 Da), additional peaks at larger m/z 801.6, 838.3, and 851.4 were visible and indicative of the AAS (**Figure 6B-C**).

**Figure 6.**
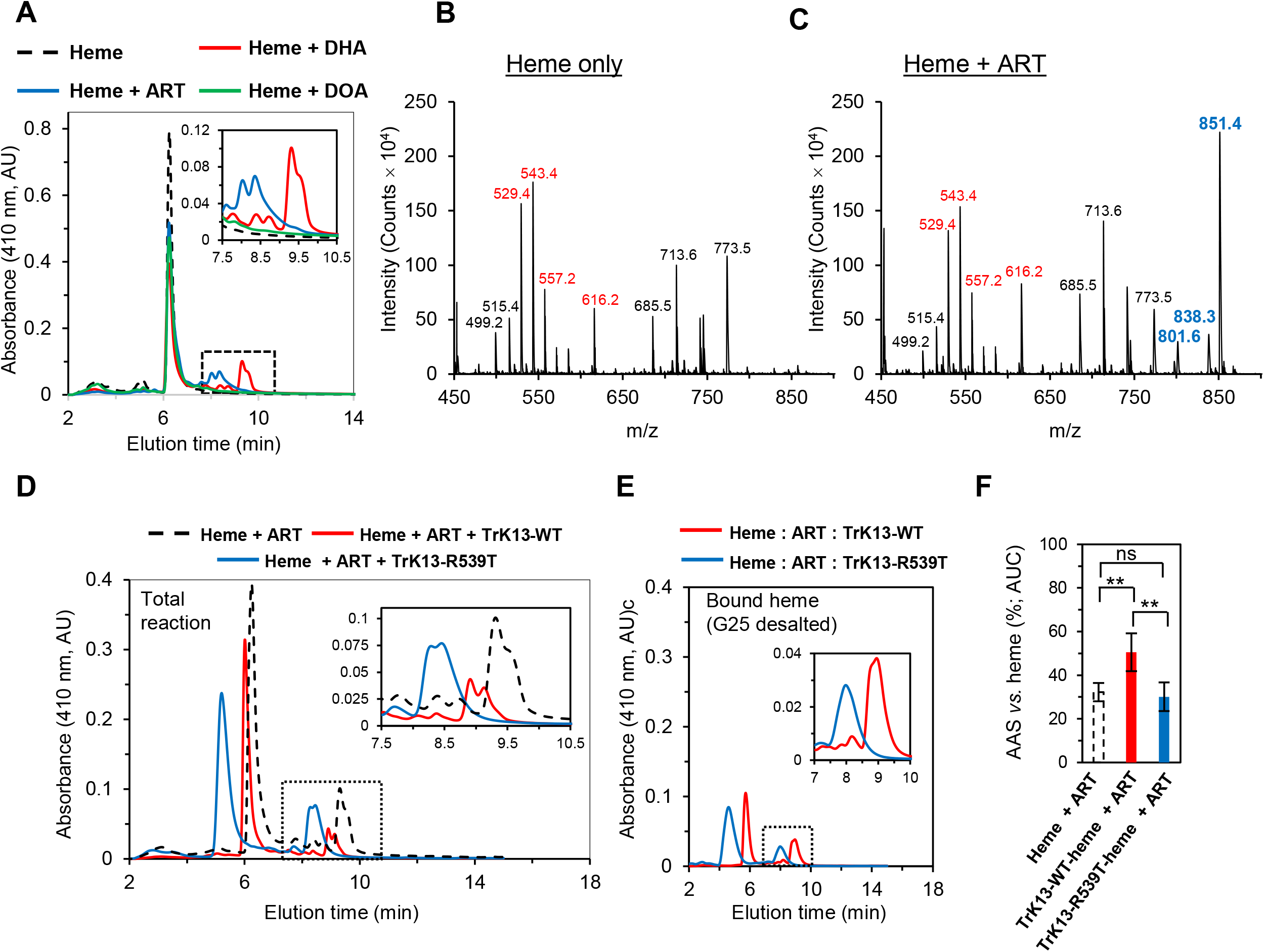
Detection of AAS by HPLC and mass spectrometry. **A.** HPLC elution profiles at 410 nm for the total reaction mixtures of heme and the corresponding peaks for free heme (tall peaks at 6-7 min) and AAS (smaller peaks at 8-10 min elution time; boxed and magnified in the inset) in the presence of either ART (red) or DHA (blue) or the inactive DOA analog (green). The elution profile of free heme is indicated by the dotted black line. **B.** MS data of the eluted samples from A (dotted box) showing the characteristic heme intensity peaks (red text) at m/z 529.4, 543.4, 616.2 and 557.2 and several other peaks generated by in-source fragmentation of heme and oxidation-reduction processes during electrospray ionization (black text). **C.** Mass spectra of the two-component mixture of heme-ART from the eluted samples of A (dotted box), showing additional AAS peaks at m/z 801.6, 838.3, and 851.4 (blue text). Peaks at these m/z were absent in the heme-only sample shown in B. **D-E.** HPLC elution profiles showing the heme (tall peaks at 5-7 min elution time) and AAS peaks (short peaks at 8-10 min elution time) in TrK13-WT (red) or TrK13-R539T (blue) containing reaction mixtures with heme (D) or in the protein-bound heme (after G-25 desalting; E). The AAS peaks are framed by dotted lines and magnified in the inset. **F.** Percentage AAS versus heme, as measured by area under the curve (AUC) for respective sample peaks and expressed as percentage. Data are mean ± SD of three independent experiments, comparison was performed by the unpaired t-test, * p < 0.05, ** p < 0.005, ns not significant (p > 0.05).

Next, we investigated whether AAS formation also occurred in the presence of TrK13-WT and TrK13-R539T proteins. TrK13 was first preincubated with heme for 60 min and then ART was added (at a molar ratio of protein: heme: ART of 1: 6: 30, *i.e*., 10: 60: 300 μM). The delayed AAS peaks were detected with both WT and mutant TrK13 proteins (**Figure 6D**), indicating that their formation was not affected by the presence of TrK13. We note that both the larger and smaller delayed peaks eluted at slightly different times depending on which variant of TrK13 was present.

The above assay lacked the ability to distinguish between the AAS formed by TrK13-bound heme or excess free heme. To address this issue, TrK13 was first incubated with equimolar concentrations of heme and desalted using a G-25 column to remove excess unbound free heme. Finally, ART was added to the desalted heme-TrK13 mixture. The resulting HPLC profile showed an overall 75% reduction in total peak heights compared to the non-desalted reaction conditions (compare peak heights between **Figure 6D** and **Figure 6E**). Interestingly, our quantitative evaluations based on the relative HPLC-resolved area-under-the-curve for the heme and AAS peak profiles suggested an enhanced AAS-forming ability of the heme molecule(s) when bound to the TrK13-WT protein (∼50%) compared to that of TrK13-R539T (∼30%) or free heme (33%) (**Figure 6F**).

We also used UV-visible spectrophotometry to quantitatively visualize and compare the ability of TrK13-bound heme to form AAS under reducing conditions^12,13^. The spectral profile of reduced heme alone remained unchanged over the 60-min recording period (**Supplementary Figure S9A**). The three combinations of two components (heme + ART, heme + TrK13, and ART + TrK13) were then tested individually as controls. When 2.5 µM heme was incubated with excess of ART (50 µM), a dramatic broadening and red shift of the heme Soret band from 400 nm to 412 nm was observed and remained stable during the 60-min experiment (**Supplementary Figure S9B**). When heme was incubated with TrK13-WT, we observed immediate hyperchromic and bathochromic/red shifts in the spectral profile of TrK13-WT, with no further changes visible during the 60-min experiment, as in our previous observations (**Figure 2D** and **Supplementary Figure 9C**). Finally, in the presence of excess ART, there were no visible changes in the spectral profile of TrK13-WT at 250-700 nm compared to the protein alone, suggesting no direct interactions between ART and the protein (**Supplementary Figure S9D**). We then prepared the heme-TrK13-WT mixture and, after desalting with G-25 column, added ART. After 30 min of incubation, we observed a shift in the absorbance spectrum across the range, with the most pronounced changes between 300-450 nm (**Figure 7A**). Specifically, we observed a hypsochromic/blue shift of the Soret peak from 410 nm to 370 nm (as previously shown in **Figure 2C**). In addition, there was a ∼42% increase in the intensity of the 370 nm peak in the presence of ART compared to heme-bound TrK13-WT without ART (**Figure 7A**). The increase in the absorbance peak at 370 nm was likely due to AAS formation, since no increase in the corresponding absorbance was detected in the control samples that were similarly treated with either the zinc-containing FPIX analog (Zn-PPIX) instead of heme (**Supplementary Figure S10**) or the inactive analog DOA instead of ART (**Figure 7B**). The formation of AAS in the same samples was then tested by LC-MS. Discrete heme peaks at m/z 512.5, 540.5, 647.5, and 664.5 were detected in the TrK13-WT-heme sample without ART, whereas additional peaks at 774.6, 801.6, and 851.4 were only detected in the sample additionally containing ART, suggesting again the formation of AAS (**Figure 7C-D**).

**Figure 7.**
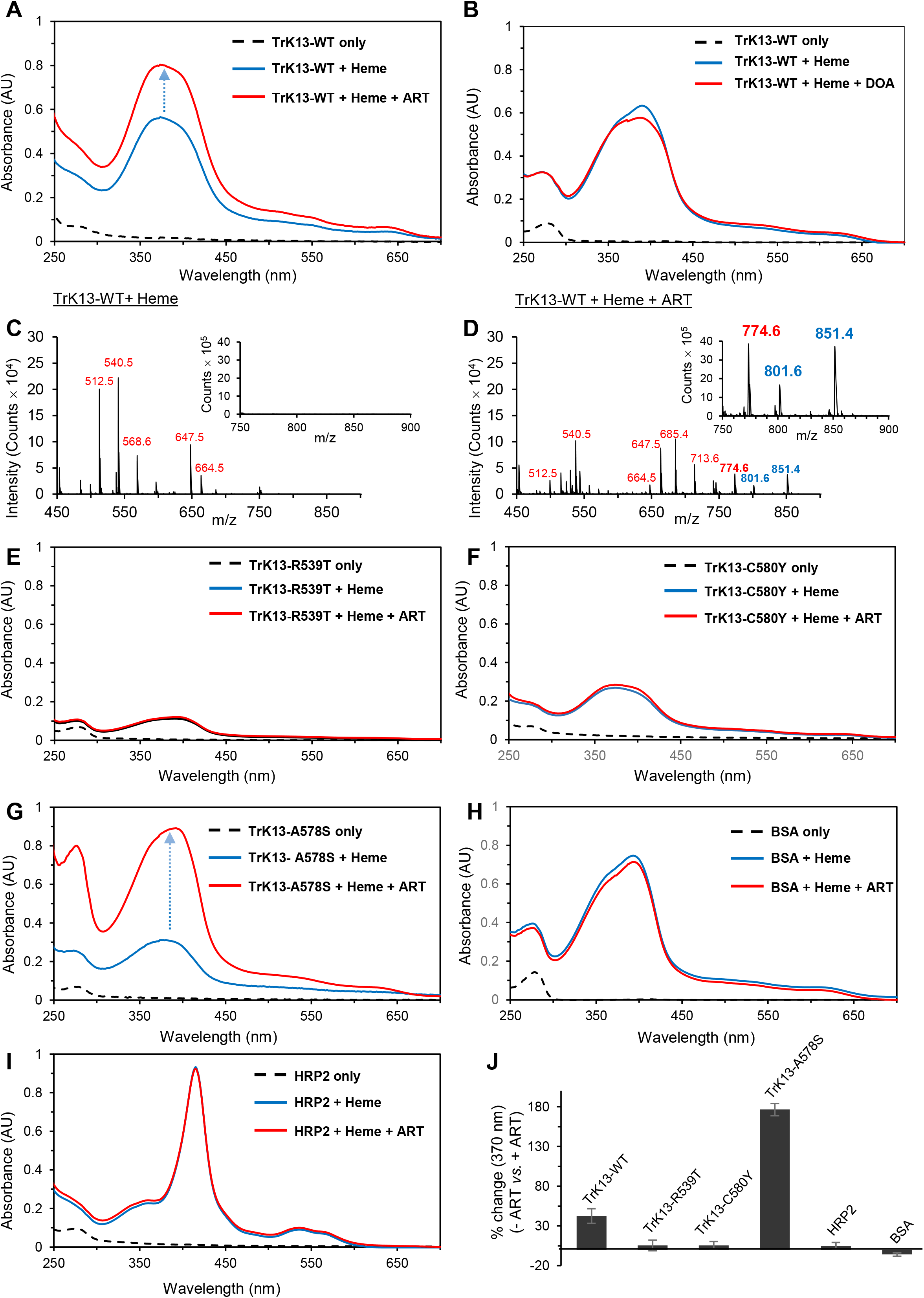
Reduced AAS formation in ART-R TrK13 mutants compared to ART-S variants. **A, E-G.** UV-visible spectroscopy scans showing further hyperchromic shift (red lines; increase indicated by blue to red gradient dotted arrow) in the absorbance profiles at 370 nm in the heme-bound ART-S TrK13-WT (A) and TrK13-A578S (G) proteins, but not in the ART-R TrK13-R539T (E) and TrK13-C580Y (F) proteins, when incubated in the presence of ART for 30 min. The Soret peaks at 370 nm (red lines) and absorption profiles of the respective proteins in the absence of heme (dotted black lines) are as indicated. **B.** UV-visible scan showing no hyperchromic shift on the Soret peak of heme-bound TrK13-WT in the presence of the inactive ART analog DOA. **C-D.** MS data of the heme-bound TrK13-WT samples (after G-25 desalting) incubated in the absence (C) or presence (D) of ART. While the peaks corresponding to heme (in red text) were visible in both samples, the peaks at m/z 801.6 and 851.4 (blue text) were only visible in the presence of ART (D) and are indicative of AAS. All samples were treated as described earlier^13^. The inset shows the expanded m/z region between 750-900. **H-I.** UV-visible spectroscopy scan showing no AAS formation by the heme-binding proteins BSA and HRP2. **J.** Graphical representation of the percentage increase in the Soret peak at 370 nm for the data in A, B, E-I Error bars are from three independent experimental replicates.

A profile similar to that of TrK13-WT was observed for the ART-sensitive TrK13-A578S protein, but with an even greater ∼176% increase in peak intensity at 370 nm (**Figure 7G**). Surprisingly, the heme-TrK13-R539T and heme-TrK13-C580Y complexes showed barely detectable increase in their respective peak intensities at 370 nm in the presence of ART (**Figure 7E-7F**), suggesting much lower AAS formation. Neither BSA (a validated heme-binding protein) nor *P. falciparum* Histidine-rich protein 2 (PfHRP2, a well-established heme-binding parasite protein^76^) showed an increase in peak absorbance at 370 nm, indicating that the ability of proteins to bind heme does not necessarily translate into AAS formation (**Figures 7H-J**; **Supplementary Figure S11**). In summary, the ART-R TrK13-R539T and TrK13-C580Y proteins exhibited reduced affinity for heme (as also observed in the MST experiments; **Figure 4**) and were associated with reduced AAS formation *in vitro,* as compared to the TrK13-WT protein.

Finally, we tested whether some of the AAS formed through the heme-TrK13 complex were released from TrK13 as soluble molecules. For this, the desalted TrK13-WT-bound heme samples were incubated in the absence (as a control) or in the presence of ART for 30 min. The reacted samples were then analyzed spectrophotometrically between 250-700 nm wavelengths both before and after a second (final) desalting step. We observed that the spectral profiles of the TrK13-heme sample incubated without ART were qualitatively similar both before and after the final desalting, showing no change in the characteristic soret peak at 370 nm (compare solid and dotted blue lines in **Figure 8A**) and essentially indicating efficient retention of heme molecule(s) with no leaching or release of bound heme during the incubation period. Interestingly, UV-visible scanning of the TrK13-WT-heme sample incubated in the presence of ART showed a dramatic decrease in the height of the 370 nm peak after the second desalting step, suggesting release and removal of low molecular weight AAS from their protein-bound state following desalting (compare solid and dotted red lines in **Figure 8A**). The height of the 370 nm peak decreased to a level even lower than that observed for the heme-TrK13-WT complex incubated in the absence of ART (compare dotted blue line *versus* dotted red line. Thus, the residual 370 nm absorbance peak in the ART-treated sample likely represented the retained heme fraction in the TrK13-WT-heme complex that was not involved in AAS formation (**Figure 8A-B**).

**Figure 8.**
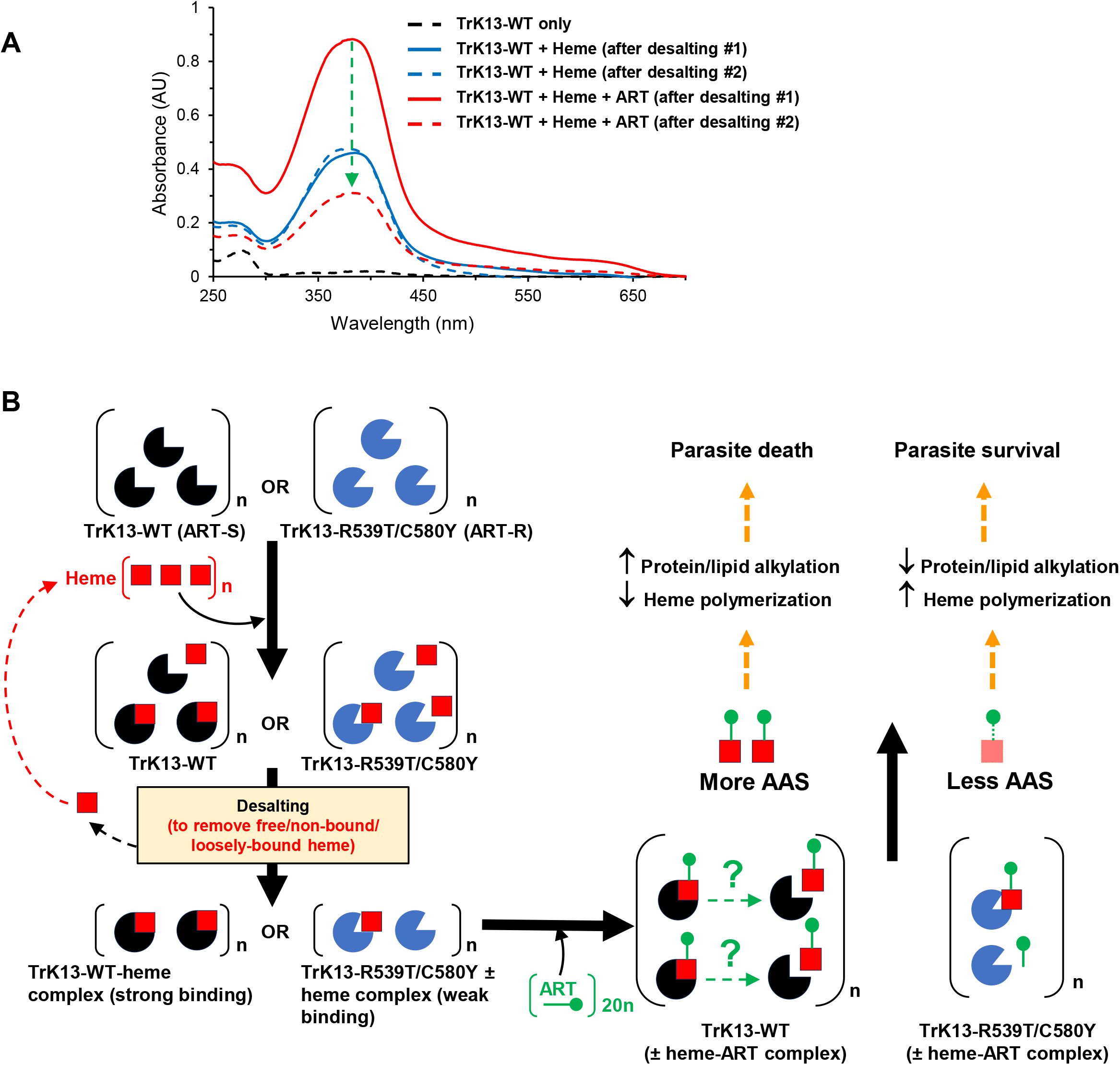
Heme bound to TrK13-WT protein is released after AAS formation. **A.** UV-visible spectroscopy scans showing the heme absorbance peak at 370 nm due to the TrK13-WT-bound heme (solid blue line) after desalting step #1 and its hyperchromic shift upon addition of ART (solid red line). The scan profile of TrK13-WT without heme is shown as a dotted black line. After desalting step #2, a steep decrease (green dotted arrow) of the heme absorbance maximum peak at 370 nm is observed in the ART-treated sample (dotted red line), indicating the release of AAS from TrK13-WT protein and its removal by the second desalting. The resultant peak at 370 nm could be due to residual bound heme that did not form AAS. No decrease in λ_max_ at 370 nm is observed in the TrK13-WT-heme-bound sample processed similarly in the absence of ART (dotted blue line), thus excluding the possibility of random heme release during the incubation period. **B.** Schematic representation of the molecules at various stages of the experiment. Recombinant ART-S (TrK13-WT; black pac-man) or ART-R (TrK13-R539T or –C580Y; blue pac-man) proteins with higher or lower heme-binding affinity (solid red square), respectively were desalted (desalting step #1) to remove the excess free heme or loosely-bound heme. Following incubation with 20-fold excess ART (green ball pin), desalting step #2 was performed to remove any AAS released from its TrK13-bound state (green question mark). Orange dotted arrow symbolizes the proposed role of liberated AAS in alkylating parasite protein targets or inhibiting hemozoin formation during the intra-erythrocytic stages of infection. Reaction mixtures at each step were analyzed by UV-visible spectrophotometry spanning 200-700 nm wavelengths.

### The native PfKelch13 protein is pulled down from *P. falciparum* lysate using hemin-agarose beads

To test the putative relevance of our *in vitro* observations with respect to the parasite-derived proteins, the RIPA-solubilized extracts from asynchronous *P. falciparum* 3D7 parasites were incubated with either hemin-agarose beads or glutathione-agarose beads (as a control) under reducing conditions. After extensive washing, the bound proteins were eluted with Laemmli buffer and resolved by SDS-PAGE. PfKelch13, was detected as a distinct ∼80-90 kDa band in western blots in the eluates from hemin-agarose beads by both the custom-generated polyclonal^47^ (**Figure 9**) and monoclonal^36^ anti-PfKelch13 antibodies (**Supplementary Figure S12**). In contrast, PfKelch13 was only faintly detected in the similarly processed samples from the control glutathione-agarose beads (**Figure 9**). Membranes were similarly probed with antibodies against either the parasite *P. falciparum* histidine-rich protein 2 (PfHRP2, a known heme-binding protein^77^; **Figure 9**) or aldolase (PfAldolase; a heme non-binding protein used as control; **Supplementary Figure S12**). Results revealed prominent PfHRP2 band in eluates from hemin-agarose beads and not in control glutathione-agarose beads. The heme non-binder PfAldolase showed background smear. Altogether, these data indicate that the native PfKelch13 was specifically recovered from the parasite solubilized-proteome through its interaction(s) with hemin-agarose beads. These results are consistent with the hypothesis that native PfKelch13 from the parasite could interact with heme.

**Figure 9.**
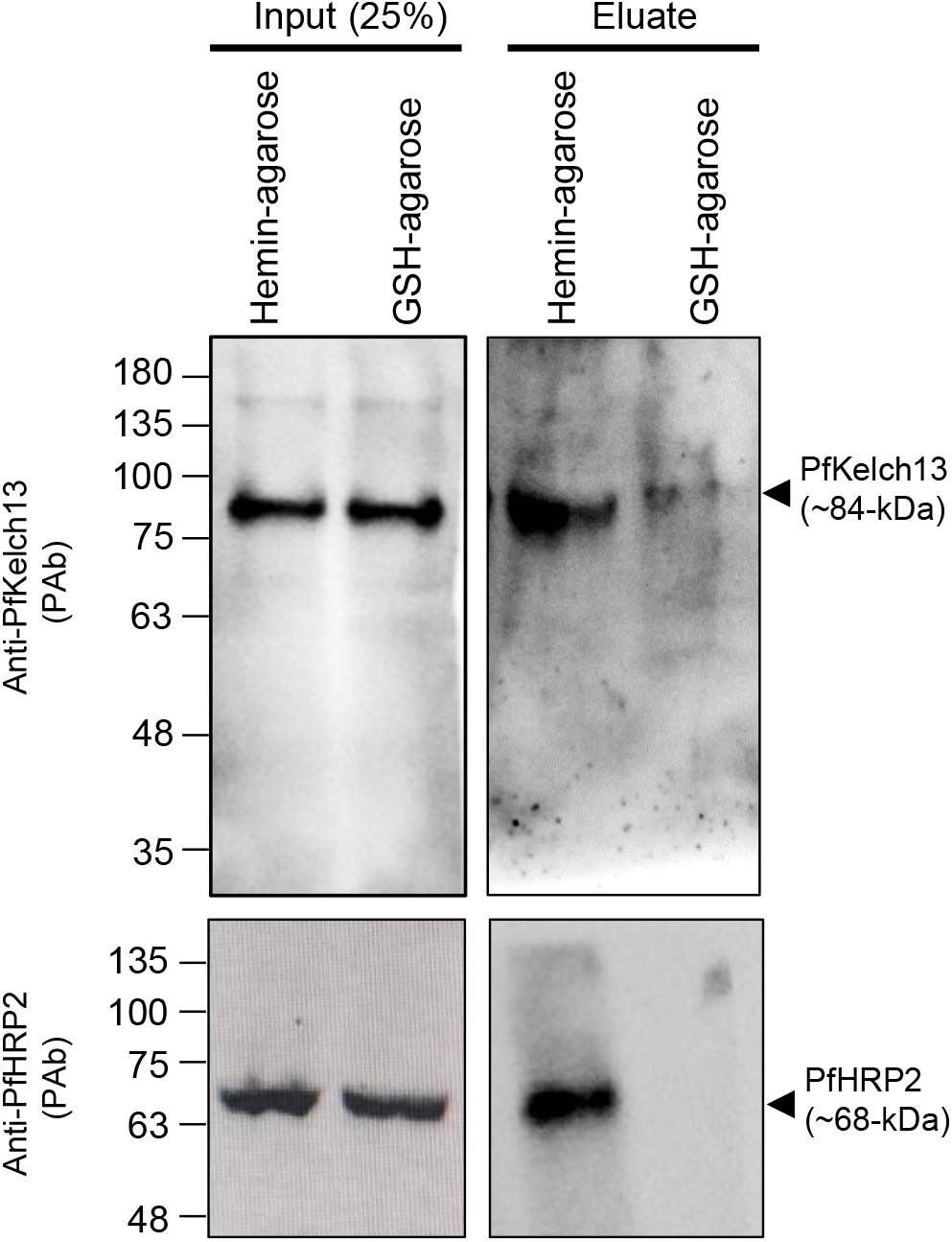
Pulldown of the native PfKelch13 protein from *P. falciparum* total protein extract with hemin-agarose beads. 3D7 parasites were isolated by saponin treatment from asynchronized culture and proteins were solubilized with RIPA buffer. Protein extracts were incubated with hemin-agarose or glutathione (GSH)-agarose beads. After extensive washing, bound proteins were eluted with Laemmli buffer, separated by SDS-PAGE, and blotted against PfKelch13 or PfHRP2 antibodies. PfKelch13 antibodies were custom-generated using a PfKelch13 peptide^47^. Molecular weight standards (in kDa) are as indicated.

## Discussion

While some cellular functions, localizations, and protein-interacting partners have been assigned to the Kelch13 protein in the apicomplexan parasites *P. falciparum*^30,36,78^ and *Toxoplasma gondii*^79,80^, unraveling the molecular consequences of ART-R mutations in PfKelch13 remains an ongoing area of investigation. PfKelch13 modulates ART sensitivity in the ring stage primarily by regulating the level of hemoglobin uptake at the parasite periphery, which translates onto decreased hemoglobin degradation into ART activator molecules^30,32^ and lower level of heme-DHA adducts^34,81,82^. The overall phenotype (reduced hemoglobin uptake and subsequent ART activation) is controlled by the decreased abundance of mutant PfKelch13 at the cytostome collar ring where hemoglobin uptake is initiated^30–32,78,84^, resulting in atypical invaginations lacking the constricted necks characteristic of cytostomes as reported in PfKelch13-mislocalized parasites^32^.

In this study, using purified N-terminally-truncated recombinant versions of PfKelch13 lacking the first 188 amino acids (TrK13)^47^, we identified *in vitro* several novel biochemical properties of PfKelch13. The purified TrK13 protein binds to iron salts and heme, an observation that was repeatedly found using a variety of techniques (**Figures 1-4**). Further, when incubated with ART, heme bound to TrK13 was associated with AAS formation. Remarkably, the binding to heme and the production of AAS were lower for the TrK13-R539T and -C580Y ART-R mutants compared to TrK13-WT and the ART-sensitive mutant -A578S proteins (**Figure 7**). Finally, the AAS formed through TrK13-WT-bound heme was likely released as soluble molecules.

Our *in silico* docking studies predicted the KRP domain of TrK13 as the major binding site for heme, carrying 7 of the 11 putative heme binding cavities for the WT (**Figure 5**). Also, the number and location of the predicted heme binding cavities varied according to the TrK13 variants, which is consistent with experimental estimates of dissociation constants (K_d_) that also varied across variants. However, which of the putative heme binding cavities in PfKelch13 are truly effective should be tested by appropriate assays.

Our comparative assessments revealed measurable differences between the heme-binding propensities of the ART-S and ART-R representative PfKelch13 proteins. The heme-mediated fluorescence quenching of the tryptophan residues was significantly more efficient in the two ART-R variants (TrK13-R539T and -C580Y) compared to the ART-S proteins (TrK13-WT and -A578S), as evidenced by the Stern-Volmer quenching constants (K_SV_)^66^, thus confirming increased accessibility (**Figure 3**). However, after careful consideration, we refrained from over-interpreting increased fluorescence quenching as increased binding affinity by these mutants for two main reasons. First, our earlier SAXS-based structural studies with the same recombinants (*i.e.*, TrK13-R539T and TrK13-C580Y) revealed detectable differences in the spatial positioning of the BTB/POZ-KRP domains of their oligomeric assemblies relative to the ART-S TrK13-WT and -A578S proteins^47^. The dimensional change in these proteins likely played an important role in their characteristic K_sv_ determination. Second, the influence of other non-binding factors such as background quenching (referred to as the inner filter effect) was also considered, particularly for heme, which absorbs both excitation light and tryptophan-emitted fluorescence to create a continuous quenching and binding effect^63,85,86^. However, the results indicated a clear pattern of difference between the ART-R and ART-S protein variants. These initial data were confirmed by subsequent MST and comparison spectra based on different dissociation constants (K_d_) in the MST experiments (**Figure 4**).

The HPLC analyses used in this study revealed distinct elution profiles of heme and AAS with expected m/z as reported previously^3,13^ (**Figure 6**). We then provided the first evidence that AAS could be successfully generated by heme molecule(s) bound to TrK13 protein(s). Unfortunately, as the amount of AAS was below the detection limit of LC-MS, we were unable to quantify the AAS formed by heme-bound TrK13-WT and TrK13-R539T, respectively. Nevertheless, the hyperchromic shift of the Soret peak of the heme-bound TrK13-WT protein in the presence of ART (by UV-visible spectrophotometry) was strongly indicative of AAS formation, as it was conspicuously absent in the case of the metabolically inactive ART derivative DOA or in the presence of ART plus the heme analog Zn-PPIX (**Figure 7**). Another interesting result of this study was the absence of detectable AAS formed by heme-BSA and heme-PfHRP2 complexes. Since BSA and PfHRP2 are two well-established heme-binding proteins^87,88, 77^, our results suggest that the heme-binding property of a protein *per se* does not translate into the propensity to form AAS (**Figure 7**). We speculate that it could be due to different heme-protein binding modes that alter the accessibility and/or chemical properties of the bound heme or iron in the heme molecule Further experiments are needed to characterize the catalytic activity of heme bound to TrK13 and to test whether it is also detected in the parasite.

In the context of the intraerythrocytic malaria parasite, there could be multiple sites of PfKelch13-heme encounters. PfKelch13 is located to various cellular compartments: at the parasite periphery, proximal or associated with the plasma membrane; close to the digestive vacuole, the apicoplast and the mitochondria; near parts of the endoplasmic reticulum (ER); and in or close to compartments associated with Rab, ATG18, and PI3P^30,36,89^. A common feature of these localization studies is that PfKelch13 was detected as cytosolic foci rather than diffuse staining, and as a single focus in early rings and multiple foci in later stages. Also, several of these cellular compartments are individually involved in hemoglobin uptake, hemoglobin degradation, and heme synthesis, respectively^30,31,35,89^. Thus, we speculate that some *de novo* or recycled free heme may either diffuse or be transported from these organelles into the parasite cytosol, where PfKelch13 is likely solvent-accessible. This is supported by the evidence of a labile heme pool (∼1.6 μM) that is stably maintained throughout the intraerythrocytic development of the parasite^42^.

Our results also indicate that some of the AAS generated by the TrK13-heme complex are likely to be soluble, as evidenced by release experiments performed with the TrK13-WT, which suggests that limited proteotoxicity of those AAS directed at TrK13 could occur. This is in line with two reports that did not identify PfKelch13 as alkylation target of artesunate or the alkyne-labeled ART analog AP1 coupled to biotin^6,9^. Nonetheless, the existence of putative TrK13-AAS adducts in our *in vitro* system should be carefully investigated by appropriate methods.

To date, the reduced abundance of PfKelch13 mutant proteins at the cellular level is considered to be an exclusive PfKelch13 feature that distinguishes ART-S and ART-R parasites^24,33,81,30^. A recent study comparatively measured the relative abundance of PfKelch13 protein in a series of PfKelch13 mutants *versus* their resistance index (RSA) within the same isogenic 3D7-derived parasites by direct fluorescence-based microscopy of N-terminus-labeled GFP-PfKelch13 variants^83^. While three PfKelch13 mutants C580Y, R539T and R561H had similar PfKelch13 abundances of 51-52% compared to the WT protein, they differed in their percentage of RSA survival (of about 16%, 38% and 11%, respectively). These results suggest that not just protein abundance but also subtle changes in PfKelch13 function due to specific mutations may modulate the level of ART-R. Results from our study also offer a plausible explanation for the previously reported ART-R to ART-S revertant phenotype in the NF54^C580Y^attB parasites (initial RSA: 4.8%) that was slightly more pronounced upon *in trans* co-expression of 3HA-PfKelch13^WT^ (RSA: 1%) than of GFP-PfKelch13^C580Y^ (RSA: 1.6%).). Although this small difference could be due to different protein levels of the co-expressed PfKelch13 (because of different tags and/or genotypes), we speculate that it could also be due to different function of PfKelch13. The *in trans* co-expression of 3HA-PfKelch13^WT^ in the NF54^C580Y^attB parasites (originally possessing a native low-affinity heme binder PfKelch13^C580Y^ protein) effectively replenished these parasites with the high-affinity PfKelch13^WT^ protein, thus enabling enhanced AAS generation and larger reversion to the ART-S phenotype.

These data obtained in the parasite are altogether in line with our *in vitro* findings that show altered biochemical properties in recombinant TrK13 mutants. We here propose that ART-R mutations in PfKelch13 would confer two distinct properties at the PfKelch13 protein level: a decreased protein abundance^32,36^ and a decreased heme-binding activity. Both properties would contribute to reduced ART-heme AAS formation but through distinct heme pools, free heme versus PfKelch13-bound heme. Since the parasite cannot reduce the hemoglobin uptake pathway too much without compromising its own survival, a direct effect of mutant PfKelch13 on reducing the production of AAS would be beneficial and complement the ART resistance phenotype associated with reduced heme supply. This scenario however assumes that heme bound to PfKelch13 would act as a better trigger for ART activation compared to cytosolic free heme, a strong hypothesis that needs to be tested experimentally. Of note, heme is also known as an important regulatory molecule in some other eukaryotic cells^90–92^. It would then be interesting to explore whether heme regulates some yet unknown, native function(s) or pathway(s) associated with PfKelch13.

In conclusion, while heme has already been validated as the ‘*spark’* for ART activation, the parasite PfKelch13 protein could serve as a double *‘valve’* to regulate the level of ART activation.

## Materials and Methods

### Reagents and chemicals

All reagents and chemicals used in this study were of analytical grade or higher and were purchased from various sources. Hemin chloride (H3741), Ferene S (82940), 3,3′,5,5′ tetramethylbenzidine (TMBZ; T0440), hemin-agarose (H6390), artemisinin (361593) and artesunate (A3731) were purchased from Merck Millipore (Germany). StrepTactin XT beads (2-5010-025) and biotin (0-1016-002) were purchased from IBA Lifesciences (Germany). Dihydroartemisinin (DHA; 19846) and deoxyartemisinin (DOA; 20426) were purchased from Cayman Chemicals (USA).

### Cloning, expression, and purification of recombinant TrK13 proteins from *E. coli*

The *E. coli* expression constructs pPR-IBA101 (PfKelch13-WT), pPR-IBA101 (PfKelch13-C580Y), pPR-IBA101 (PfKelch13-R539T) and pPR-IBA101 (PfKelch13-A578S) were generated as described in our previous study^47^. The recombinant proteins TrK13-WT, TrK13-C580Y, TrK13-R539T and Trk13-A578S with a C-terminus-fused strep tag were purified from isopropyl-1-thio-β-D-galactopyranoside (IPTG) and arabinose (0.2%) induced culture lysate using StrepTactin XT Superflow resin (IBA, Germany). The purified proteins were dialyzed overnight at 4°C against MOPS buffer, pH 7.4 with at least three periodic changes. Protein samples were then concentrated using Amicon 3-kDa concentrators (Millipore), and the purity of each protein was assessed by SDS-PAGE and further confirmed by Western blotting using commercial antibodies against the strep tag (Biobharati Life Sciences) or custom-generated anti-PfKelch13^47^.

The pPR-IBA101 (PfKelch13-QNG) plasmid was generated using an overlapping PCR strategy. Briefly, the pPR-IBA101 (PfKelch13-WT) plasmid was used as a template for two independent PCRs (PCR1 and PCR2) with PfKelch13-BsaIF (5’-CAAATGGGAGACCTTATGGAAGGAGAAAAAGTAAAAACAAAAGCAAATAGTATCTCG-3’)/ K13-EEHtoQNG-R (5’-AAAGAATCCACATGAATTTAGAACATTGCCATTTTGTCCTCCTGTAATTATATAAGAATCTGACAAT GTGGC-3’) and K13-EEHtoQNG-F (5’-GGAGGACAAAATGGCAATGTTCTAAATTCATGTGGATTCTTTTCACCAGATACAAATGAATGGC-3’)/PfKelch13-BsaIR (5’-CCAAGCGCTGAGACCAGCAGCTATATTTGCTATTAAAACGGAGTGACCAAATCTGGG-3’) primer pairs. These PCR products were used as templates in PCR3 to reconstitute *pfkelch13-qng*, which was then digested with BsaI and cloned into the appropriate site in pPR-IBA101. The resulting E688Q, E691N and H697G mutations in pPR-IBA101 (PfKelch13-QNG) were confirmed by Sanger sequencing and the recombinant protein TrK13-QNG was purified in a similar manner.

The pPR-IBA101 (PfHRP2) plasmid was prepared by amplifying *pfhrp2* (total length 1064 base pairs) from nucleotide positions 228-1061 (excluding the signal sequence coding region) by PCR using *P. falciparum* genomic DNA as template and the primers NoSS-HRPII-BsaIF (5’-CAAATGGGAGACCTTGCATTTAATAATAACTTGTGTAGCAAAAATGCAAAAGG-3’) and HRPII-NS-BsaIR (5’-CCAAGCGCTGAGACCAGCAGCATGGCGTAGGCAATGTGTGG-3’). The PCR product was then digested with BsaI and cloned into the appropriate site in pPR-IBA101 and confirmed by Sanger sequencing. The recombinant PfHRP2 was purified as described above for TrK13 proteins.

### *In silico* prediction of Fe^2+^ and Fe^3+^ binding sites in full length PfKelch13 and TrK13-WT with C-terminal strep tag fusion

The binding sites for Fe^2+^ (ferrous) and Fe^3+^ (ferric) were predicted using the MIB server (http://bioinfo.cmu.edu.tw/MIB/)^50,93^. MIB is a metal ion binding site prediction and docking server that provides an integrated approach to searching for residues involved in metal ion-binding sites using the fragment transformation method. The PfKelch13 amino acid sequence was compared with Fe^2+^ (ferrous) and Fe^3+^ (ferric) metal-binding templates in the database to locate putative metal-binding residues. Each residue of the query protein is assigned a binding score, which is a combination of sequence and structure conservation measures. If the binding score of a residue is higher than a specified threshold, that residue is predicted to be a metal-binding residue. Based on the local 3D structure alignment between the query protein and the metal ion-binding template, the metal ion in the metal-binding template can be transformed into the structure of the query protein.

### UV-visible spectrophotometry-based detection of interactions between recombinant TrK13-WT protein and metal salts

The absorbance and peak profile of purified recombinant TrK13-WT protein in the absence or presence of excess metal salts were measured at a wavelength of 250-700 nm using a JASCO UV600 spectrophotometer with a path length of 1 cm, and changes therein were considered indicative of protein-ligand interactions. Briefly, 500 µM of individual salt solutions of different metals [ferrous ammonium sulfate (FAS), ferric ammonium sulfate (FES), magnesium sulfate (MgSO_4_), manganese chloride (MnCl_2_), zinc chloride (ZnCl_2_), and copper sulfate (CuSO_4_)] in 50 mM MOPS buffer, pH 7.4 were scanned at a wavelength of 250-700 nm. As a control, purified recombinant TrK13-WT protein (5 µM in 50 mM MOPS buffer, pH 7.4) was also scanned at these wavelengths. Subsequently, reaction mixtures containing 5 µM TrK13-WT (or mutants) and 500 µM metal salts were incubated for 30 min and then scanned. Experiments with TrK13 mutants were performed similarly, using FAS and FES only.

### Estimation of iron-binding by Ferene S colorimetric assay

Ferene S [3-(2-pyridyl)-5,6-bis (2-(5-furylsulfonic acid))-1,2,4-triazine, disodium salt] colorimetric assay was used according to published protocol^94^ as a specific stain to detect iron molecules bound to the recombinant TrK13-WT protein. All glassware used in the assay was previously soaked in 1% (v/v) HCl to remove iron ions. Purified recombinant TrK13-WT protein (5 μM) was incubated for 60 min with 450 μM FAS in 50 mM MOPS buffer. As control, a sample with 450 μM FAS only (without TrK13) in 50 mM MOPS buffer was processed similarly. After incubation, free unbound salt was removed by size-exclusion chromatography using a PD Spintrap™ G-25 desalting column and the eluate containing biomolecules larger than 5 kDa was incubated with 30 μl of 10 M HCl for 10 min under rotating conditions. Then 25 μl of 80% trichloroacetic acid (TCA) was added and kept on ice for 10 min. The solution was then centrifuged at 10,000 × g for 10 min and the supernatant was transferred to a cuvette. Then, 30 μl of 45% acetic acid and 100 μl of freshly prepared Ferene S reagent (containing 45% w/v sodium acetate, 10 mM ascorbic acid, and 0.75 mM Ferene S reagent) were added and the sample was immediately mixed. The absorbance at 595 nm was then measured. In this assay, iron-containing solutions turn blue.

### Preparation of Fe-NTA beads and pull-down assays using recombinant TrK13-WT protein

Fe-NTA agarose beads were prepared from commercially available Ni-NTA beads (Qiagen) according to previously described protocols^53,54^. Briefly, 1 ml of a 50% v/v suspension of Ni-NTA was washed with 20 ml of 100 mM EDTA to remove any bound Ni^2+^ ions. The beads were then sequentially washed with 30 ml of double-distilled water, 30 ml of 0.1 M acetic acid, and 30 ml of 100 mM iron salts (FAS, FES, or FeSO4) to load with the appropriate metal ions. After binding, the beads were washed again with 30 mL of water followed by 30 ml of 0.1 M acetic acid to remove any unbound metal ions. Finally, the Fe-NTA beads were equilibrated with buffer W (10 mM Tris/HCl pH 8.0, 1 mM EDTA) and used for pull-down assays. The loading conditions assumed 100% replacement of Ni^2+^ by iron salts. For pull-down assays, 5 µM affinity-purified TrK13-WT or TrK13-R539T protein were co-incubated with Fe-NTA or Ni-NTA beads in buffer W in the absence or presence of excess (0.1 or 10 mM) free FAS or FES salts for 1 h at room temperature. The beads were thoroughly washed with buffer W, and Fe-NTA-bound proteins were resolved by SDS-PAGE followed by Coomassie staining.

To analyze the effect of TrK13-WT oligomerization status on Fe-NTA agarose binding, the affinity-purified protein (in 50 mM MOPS buffer, pH 7.4) was incubated either without or with increasing concentrations of urea (0-6 M) for 30 min, followed by cross-linking with 0.1% (v/v) glutaraldehyde (with spacer length of 5 Å). The samples were then incubated with 30 µl of 50% v/v resuspended Fe-NTA beads at RT for 1 h, washed with the same buffer to remove unbound proteins, resolved by SDS-PAGE and stained with Coomassie.

### Sequence and structural overlap between the KRP domains of TaTFP and PfKelch13

Pairwise alignments between the amino acids of the KRP domains of TaTFP and PfKelch13 were performed using Clustal Omega (https://www.ebi.ac.uk/Tools/msa/clustalo). The PDB structures of TaTFP (RCSB PDB ID: 5A11) and PfKelch13 (RCSB PDB ID: 4YY8) were retrieved from the RCSB database (https://www.rcsb.org) and aligned using PyMOL software (https://pymol.org).

### *In gel* staining with TMBZ to detect heme-binding to the TrK13-WT protein

*In gel* staining of TrK13-WT protein with 3, 3′, 5, 5′ tetramethylbenzidine (TMBZ) was performed according to a published protocol^55,56^. Briefly, 10 µM of affinity-purified recombinant TrK13-WT was preincubated in the absence or presence of an equimolar (10 µM) concentration of heme and resolved by PAGE under native conditions. The gel was incubated for 2 h in the dark with freshly prepared TMBZ solution (6.3 mM TMBZ was dissolved in methanol and mixed with 0.25 mM sodium acetate pH 5.0 in a 3:7 ratio). The bands were developed by adding 30 mM H_2_O_2_ for 30 min. Finally, the gel was fixed by three washes with isopropanol and 0.25 mM sodium acetate pH 5.0 in a 3:7 ratio.

### UV-visible spectrophotometry for salt and heme-binding to the recombinant proteins

All spectrophotometric experiments were performed in an 8-well microcuvette using a JASCO UV660 spectrophotometer with 1 cm path length. The total reaction volume was 100 µl in all cases. For salt-binding assays, 5 µM concentrations of affinity-purified recombinant proteins (TrK13-WT or the corresponding -C580Y, -R539T, and -A578S) were incubated with 100-fold excess (500 µM) concentrations of FAS, FES, MgSO_4_, MnCl_2_, ZnCl_2_, or CuSO_4_. Reaction mixtures containing the protein and/or the respective salts were prepared in MOPS buffer, pH 7.4. After incubation for 1 h, the samples were scanned between 250-700 nm.

For heme-binding assays, a stock solution of reduced heme was prepared by dissolving hemin chloride in 0.1 M NaOH, followed by dilution to the respective working concentrations in MOPS buffer, pH 7.4, in the presence of 2 mM sodium dithionite (to reduce hemin to heme). Affinity-purified recombinant TrK13 proteins (WT, C580Y, R539T and C580Y) or commercially purchased BSA (A7030) or lysozyme (L6876) (Merck, Germany), each at 10 µM concentration, were incubated in the absence or presence of an equimolar concentration of reduced heme for 1 h at room temperature (RT) and immediately desalted using PD Spintrap™ G-25 columns (Cytiva) to remove excess unbound heme. The eluted fractions were then analyzed by spectrophotometric scanning between 250-700 nm wavelengths. Protein samples incubated in the absence of heme were processed similarly and served as appropriate controls. Experiments with the heme analogs PPIX and Zn-PPIX were also performed according to the same protocols.

To calculate the dissociation constant (K_d_) for heme binding by the TrK13-WT protein, increasing concentrations (0-20 µM) of the recombinant protein were incubated with a fixed concentration (10 µM) of heme and processed similarly. The values for the absorbance maxima at 370 nm (y-axis) were plotted against the respective protein concentrations (x-axis) as previously described^95^. In all cases, the composite data were plotted using Excel software.

### Binding of recombinant TrK13 proteins to hemin-agarose beads and competitive inhibition by excess heme and analogs

The affinity-purified recombinant TrK13 proteins (10 µM) were incubated with 30 µl suspensions of 50% v/v hemin-agarose beads for 1 h at RT in MOPS buffer, pH 7.4, under rotating conditions, as described elsewhere^96,97^. Excess unbound proteins were removed by successive five washes with ten volumes of MOPS buffer, pH 7.4, and bound fractions were solubilized in Laemmli’s sample buffer, resolved by SDS-PAGE, and visualized by Coomassie staining. For competitive inhibition, assays were performed under similar conditions in the absence and presence of excess heme or heme analogs. Briefly, 10 µM TrK13-WT protein was preincubated with 10-fold molar excess (100 µM) concentrations of reduced heme, PPIX, or Zn-PPIX for 1 h at RT, followed by the addition of hemin-agarose beads. Percentage inhibition was calculated based on the band intensity of hemin-agarose bound TrK13-WT protein without any heme or analogs and considered as 100%. The pixel intensity of each band was calculated and plotted using ImageJ software.

### Tryptophan fluorescence quenching

Tryptophan fluorescence quenching experiments were performed using an FLS920 fluorescence spectrophotometer (Edinburgh Instruments, UK). Briefly, 10 µl of TrK13-WT protein or corresponding mutants were incubated with stepwise increasing (0-5 µM) concentrations of heme as a quencher. The excitation wavelength was 295 nm and the emission wavelength was 300-500 nm. The fluorescence quenching constant (K_SV_) was calculated from the Stern-Volmer equation as described in Edgar *et al.* (2016)^56^; F_0_/F= 1 + k_q_ζ_0_[Q]) 1 + K_SV_[Q](1), where F_0_ and F are the fluorescence intensities before and after quencher addition, respectively, k_q_ is the bimolecular quenching constant, ζ_0_ is the lifetime of the fluorophore in the absence of quencher, and [Q] is the concentration of quencher. The dissociation constant (K_D_) was calculated from the change in fluorescence intensity (ΔF) at 334 nm and the concentration of heme using SigmaPlot v12.3 and the following equation: ΔF = (ΔFmax × [heme])/ (K_D_ + [heme]).

### Circular dichroism (CD) of TrK13-WT protein and heme

CD data of recombinant TrK13-WT protein were collected in the absence or presence of heme at concentrations of 0-40 µM using an AVIV circular dichroism spectrophotometer (model 420SF, Lakewood, NJ, USA). Spectra were recorded in a 0.2 cm path length quartz cuvette with a bandwidth of 1.0 nm and a resolution of 200-250 nm. A total of five scans were recorded and then averaged to obtain a better signal-to-noise ratio. Experiments were performed in a selection buffer (150 mM NaCl and 20 mM Tris). HCl (pH 8) in a concentration range of 2-5 µM protein at a scan speed of 20 nm/min. The data were processed to estimate the secondary structure content of the protein using the K2D2 server^68^.

### Microscale Thermophoresis (MST)

The binding affinity of each recombinant TrK13 protein for heme was evaluated by MST analysis using the Monolith NT.115 instrument (NanoTemper Technologies). The recombinant TrK13 proteins (50 nM) were diluted in phosphate-buffered saline (PBS; pH 7.4) supplemented with 0.05% Tween 20 (PBS-T) to prevent sample aggregation, followed by labeling with cysteine-reactive dye (30 mM; Monolith Protein Labelling Kit Red-Maleimide 2nd Generation, NanoTemper) and incubated for 30 min at room temperature (RT) in the dark. The labeled PfKelch13 protein and buffer were added to an equilibrated column and the elution fractions were collected. Elutes with less than 1,000 fluorescence counts were used for interaction analysis. Labeled TrK13-WT was titrated with hemin chloride (0.01 nM to 1 mM). Samples were premixed and incubated for 10 min at RT in the dark before loading onto standard treated capillaries (K002 Monolith NT.115). For interaction analysis, the change in thermophoresis was expressed as the fluorescence change in the MST signal, defined as F_hot_/F_cold_ (F_hot_ as the hot region after IR laser heating and F_cold_ as the cold region at 0 s). Titration of the non-fluorescent ligand results in a gradual change in thermophoresis, which was plotted as DF_norm_ to obtain a dose-response (binding) curve that can be fitted to derive binding constants. Data evaluation was performed using Monolith software (Nano Temper). Measurements were performed using standard capillaries from 25°C at 40% MST power. The thermographs showed no aggregation or molecular adsorption to the capillaries (data not shown). The smooth binding curve relating heme concentration to normalized fluorescence was fitted using a K_d_ model. Each point represents the average of three sets of individual measurements.

### *In silico* docking of the PfKelch13 protein structure with heme

The CB-Dock2 web server was used to search for cavities and heme-binding interaction sites in the PfK13 protein^98^. Briefly, CB-Dock2 used a protein surface curvature-based cavity detection approach to guide molecular docking with AutoDock Vina^99^. The TrK13-WT hexameric structure, one monomer of TrK13-WT, TrK13-R539T, TrK13-A578S, and TrK13-C580Y, and one monomer of the 1.81 Å resolution X-ray structure of WT PfKelch13 BTB-KRP (PDB ID: 4YY8) were each submitted to CB-Dock2 with the heme molecule in PDB format (retrieved from https://www.chemspider.com/Chemical-Structure.4802.html). We searched for a maximum of 20 cavity numbers. All detected cavities were used to measure the binding free energy with heme. The binding free energies were ranked according to the Vina score (in kcal/mol). The lower the value, the stronger the interaction between K13 and heme. The docking results were visualized in PyMOL Molecular Graphics System (Schrödinger, LLC).

### Detection of AAS by High-Performance Liquid Chromatography (HPLC)

HPLC analyses were performed using a GSK Gel Protein 5 µm C4-300 4.6 × 150 mm semi-preparative column and a flow rate of 1 mL/min by modification of previously described methods^100^. The reaction mixture containing 300 µM heme in the absence or presence of 360 µM ART or ART derivatives (DHA, AS) or the metabolically inactive analog DOA (1:1.2 molar ratio) was incubated for 30 min at 37°C under reducing conditions (2 mM sodium dithionate). The reaction products were separated by HPLC using the following order of eluents: (A) 0.05% v/v trifluoroacetic acid in water, (B) 0.05% v/v trifluoroacetic acid in acetonitrile. The elution gradient was as follows: from A/B=63/37 to A/B=52/48 in 15 min, then A/B=52/48 for 15 min. The flow rate was 1 mL/min and the UV detection was set at 410 nm for heme detection (Soret peak).

For assays containing the purified TrK13 proteins, 10 µM TrK13 were incubated with 60 µM reduced heme for 30 min at 37°C, followed by the addition of 72 µM ART. The reaction mixtures were incubated again for 30 min, loaded onto the HPLC column and eluted as described above. In some cases, the heme-TrK13 mixtures were desalted using G-25 columns after the 30 min incubation to remove unbound heme and before the addition of ART.

### Identification of AAS by Liquid Chromatography-Mass Spectrometry (LC-MS)

Reaction mixtures containing heme incubated in the absence or presence of ART were extracted with 1.5 ml CHCl_3_: H_2_O (2:1) as described elsewhere^13^. The organic layer was separated, dried under vacuum, and redissolved in MS grade methanol and 0.1% formic acid for LC-MS analysis. Data were collected in positive ion mode using an Agilent Q-TOF G6550A mass spectrometer equipped with a dual ASI ion source. The electrospray ionization source was operated at a nozzle voltage of 0.5 kV with N2 as the nebulizer gas. The m/z acquisition range was 100-1000 and the scan rate was 1 spectra/sec. A similar protocol was followed with slight modifications for the LC-MS analysis of samples containing TrK13. Briefly, 10 µM TrK13-WT was pre-incubated with 60 µM heme and PD Spintrap™ G-25 columns were desalted to remove free unbound heme, followed by the addition of 72 µM ART.

### Detection of AAS by UV-visible spectrophotometry

Ten micromolar purified TrK13 or commercial BSA (A7030, Merck, Germany) was incubated in the absence or presence of an equimolar concentration of reduced heme or the heme analog Zn-PPIX for 1 h at room temperature and immediately desalted using PD Spintrap™ G-25 columns (Cytiva) to remove excess unbound heme according to the manufacturer’s instructions. The G-25 column eluate, containing biomolecules larger than 5 kDa (*i.e.,* TrK13 and heme-TrK13 complex), were then incubated with 200 µM ART or DOA for 30 min and analyzed by spectrophotometric scanning between 250-700 nm wavelengths to detect the presence of a Soret peak from heme or Zn-PPIX. Control samples without heme and/or without ART were included when appropriate.

To determine whether the AAS were released from the TrK13 protein (or from other control proteins), we followed the same protocol as above except that the final reaction mixture was desalted using PD Spintrap™ G-25 columns and the eluate containing low molecular weight compounds was analyzed by spectrophotometric scanning between 250-700 nm wavelengths.

### Parasite culture and pulldown assays

*P. falciparum* 3D7 parasites were grown in human erythrocytes supplemented with 2 mM L-glutamine, 50 mg/l hypoxanthine, 25 mM HEPES, 0.21% NaHCO_3_, 0.1 mg/l penicillin-streptomycin, and 0.5% w/v Albumax II (Invitrogen) under 5% CO_2_ and 5% O_2_, at 37°C. Approximately 2 × 10^9^ 3D7 parasites were isolated by saponin treatment from asynchronized culture (4 flasks of 25 mL culture at 4% parasitemia) and frozen immediately at -80°C as dry pellets (total parasite pellet volume was approximately 100 µl). Total parasite proteins were solubilized with RIPA buffer as follows: 1 volume of dry parasite pellet was resuspended in 10 volumes of ice-cold RIPA buffer complemented with 1X protease inhibitor (cOmplete, Roche). After one hour of incubation on ice with regular mixing, the sample was passed 4 times through a 21G needle. After centrifugation (15 min, 21,100 × g, 4°C) the supernatant containing soluble proteins was collected. 48% of this soluble protein extract was incubated with 35 µl of hemin-agarose or glutathione (GSH)-agarose bead pellets that had been pre-washed 3 times with 1X PBS and once with RIPA buffer. The remaining 4% of protein extract were kept frozen at -80°C. The protein extract/beads mixture was incubated overnight at +4°C on a rotating wheel and then centrifuged (5 min, 500 × g, 4°C). The supernatant was stored as flow through and the beads were washed 6 times with 1 ml RIPA buffer complemented with 0.25× protease inhibitor (cOmplete, Roche). Bound proteins were eluted from agarose beads with 1 bead volume of Laemmli buffer heated at 95°C for 5 min. After a brief centrifugation (30 sec, 2 000 × g, room temperature), proteins were electrophoresed on precast 4–20% Tris-Glycine gels (Bio-Rad) and transferred onto nitrocellulose membranes which were blotted against PfKelch13 (antibody 1: 1/1000, E9 mouse monoclonal antibody raised against the PfKelch13 propeller domain and generously provided by Prof. David Fidock^36^; antibody 2: generated in rabbits against the peptide CEHRKRFDEERLRFL corresponding to amino acids 297-310 of PfKelch13 sequence^47^, PfAldolase (1:1000, Ab207494, Abcam), and PfHRP2^101^ antibodies. Anti-mouse (1:5000 (StarBright Blue 700, Bio-Rad) and anti-rabbit (1:5000, StarBright Blue 520, Bio-Rad) secondary antibodies were used and western blots were imaged on a ChemiDoc system (Bio-Rad). Replicate experiments were conducted in parallel and independently.

## Author contributions

Conceptualization Ideas: F.A, P.C., R.C., J.C. and S.B.

Methodology: A.R., S.T., R.C., N.K., S.D., M.F., J.R.N. and H.R.

Validation: A.R., S.T., R.C., N.K., S.D., M.F., J.R.N. and H.R.

Investigation: R.C., P.C., J.C. and S.B.

Resources: R.C., P.C. and S.B.

Writing: R.C., J.C. and S.B.

Supervision: S.B.

Project Administration: S.B.

Funding acquisition: S.B. and J.C.

## Acknowledgements

The authors would like to thank Dr. Samir K. Nath, Bhumika Vaidya and the Proteomics facility CSIR-Institute of Microbial Technology, Chandigarh, India for help in the HPLC studies, sample processing and LC-MS analyses. The authors also acknowledge the help from the Protein Structure facility at Indian Institute of Technology, Delhi (IIT-D) for the fluorescence quenching and MST studies. This work was supported by funding from the Indian Council of Medical Research (ICMR), New Delhi (58/25/2020/PHA/BMS) and partly by the Agence Nationale de la Recherche (ANR-17-CE15-0013-03 to J.C). A.R., S.T., H.R. are recipients of research fellowship from the Indian Council of Medical Research (ICMR) and University Grants Commission/Council of Scientific and Industrial Research, Government of India and IIT-Delhi, respectively. J.R.N. is the recipient of DST INSPIRE Fellowship. We thank Prof. David Fidock for the generous gift of the monoclonal PfKelch13 antibody. We would also like to acknowledge Saji Menon, Senior Field Application Scientist and Ruchika Dadhich, Field Application Specialist, NanoTemper Technologies, GmbH for carrying out the MST studies.

## Funding sources and disclosure of conflicts of interest

The authors declare that the funding source(s) had no involvement in the design of this study, data collection, analyses and interpretation and in the decision to submit the article for publication.

The authors also declare no conflicts of interest associated with this manuscript.

## Supplementary data

Supplementary Tables 1 and 2

Supplementary Figures S1-S12

**Supplementary Table 1.**
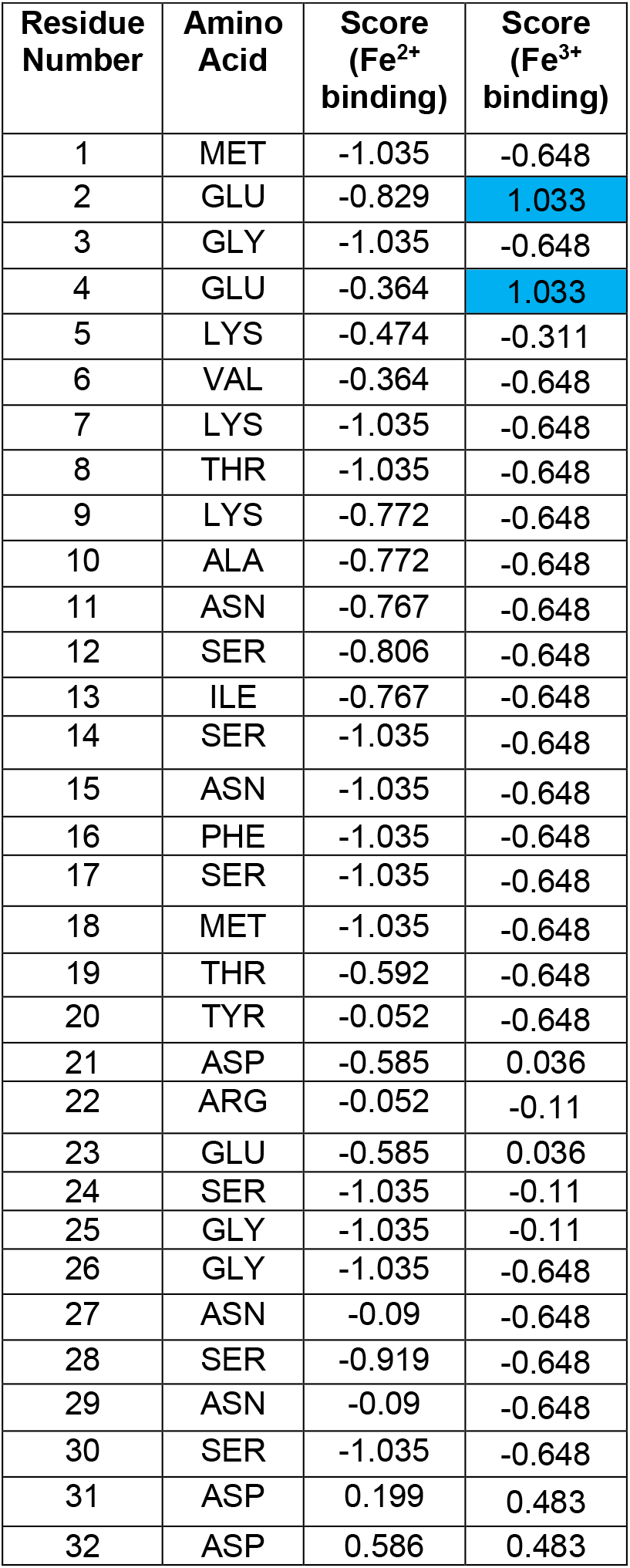
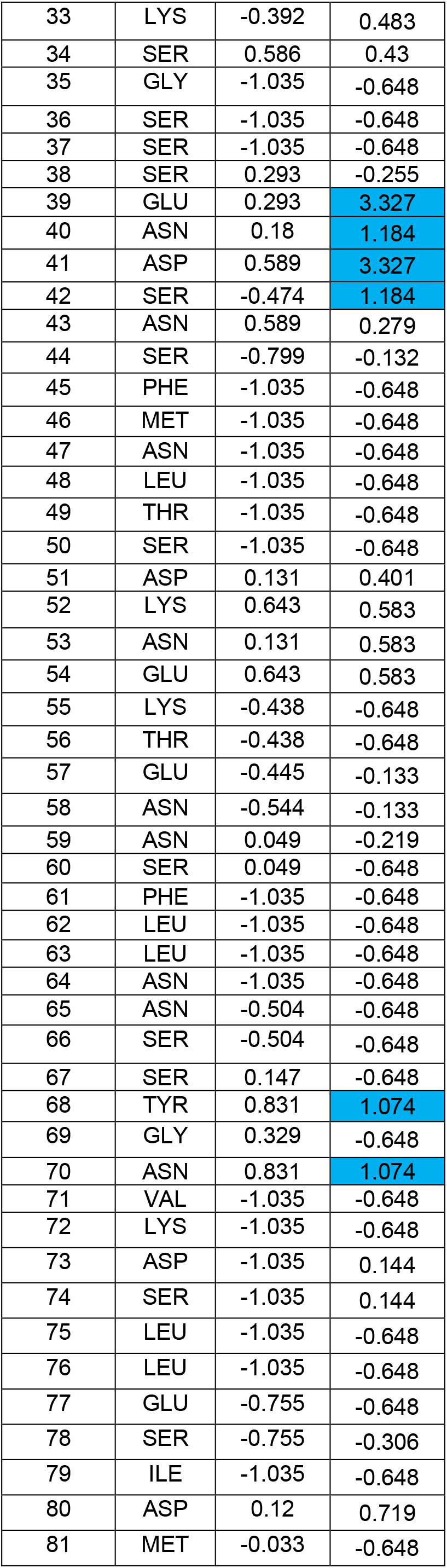
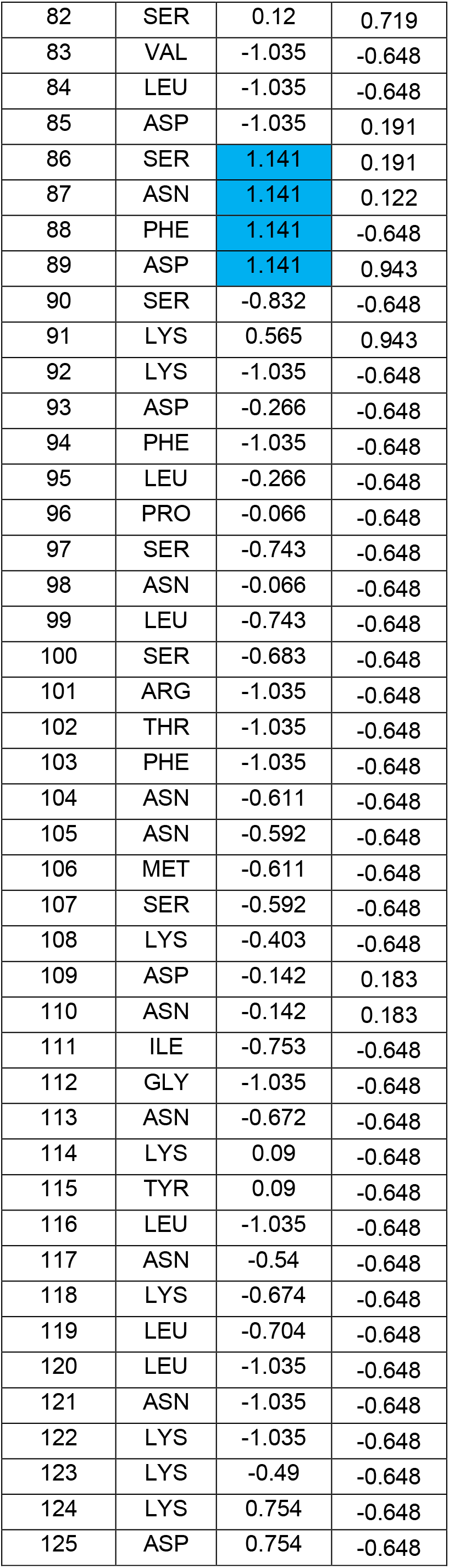
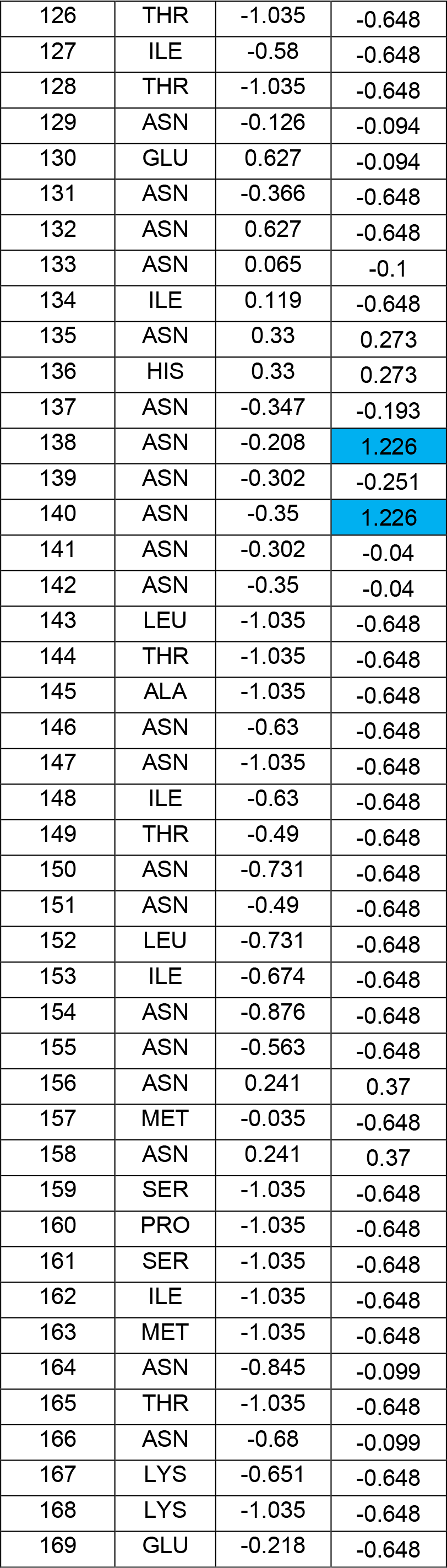
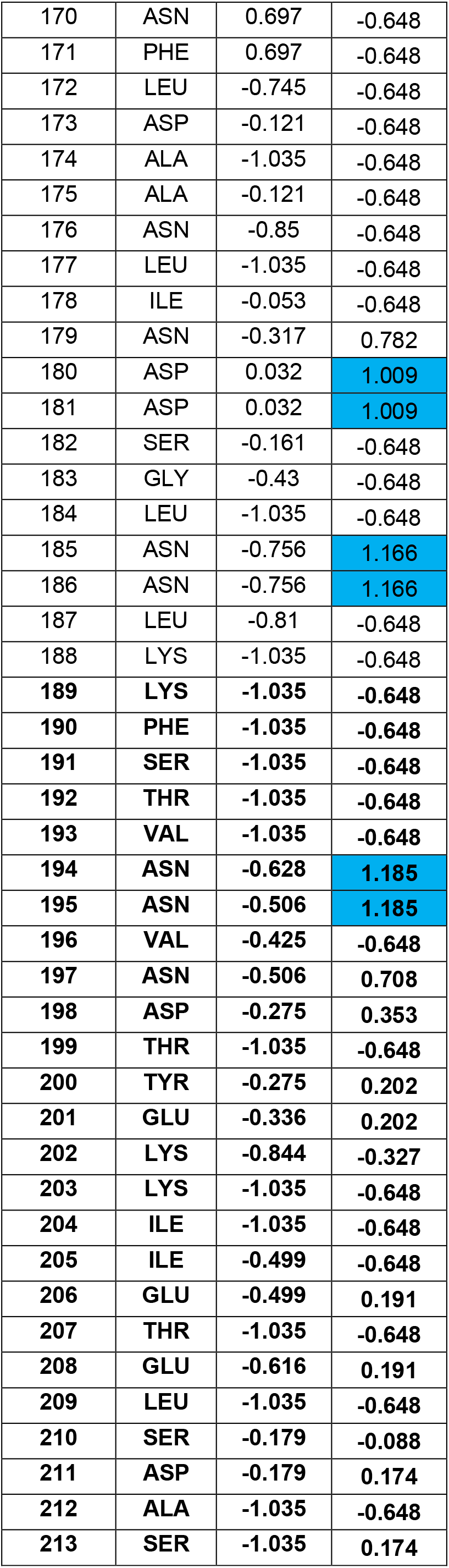
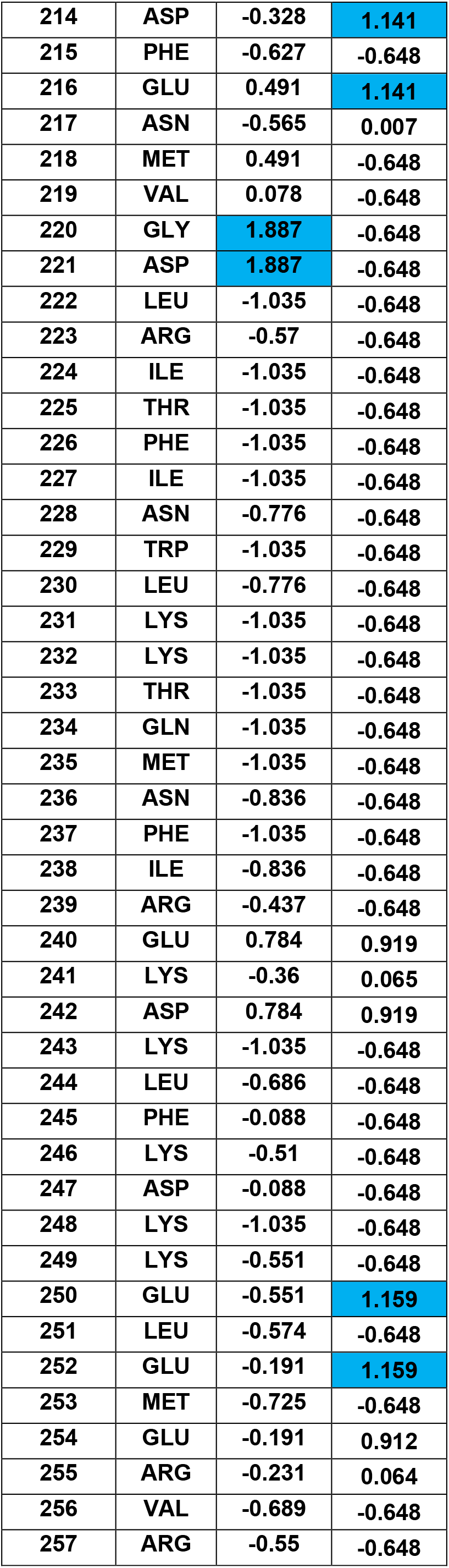
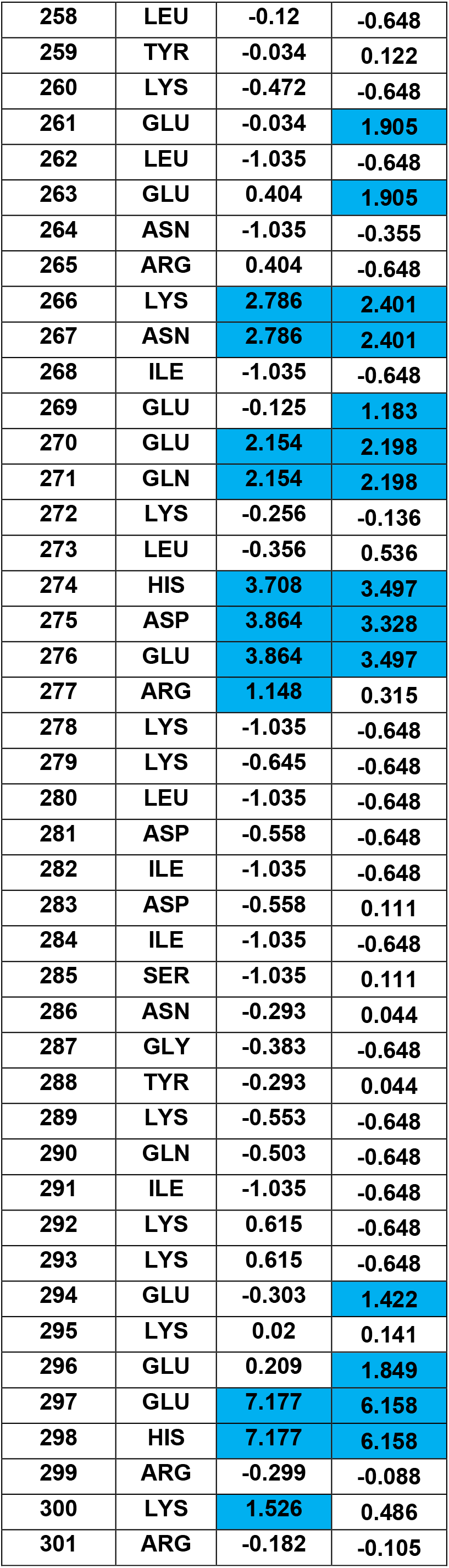
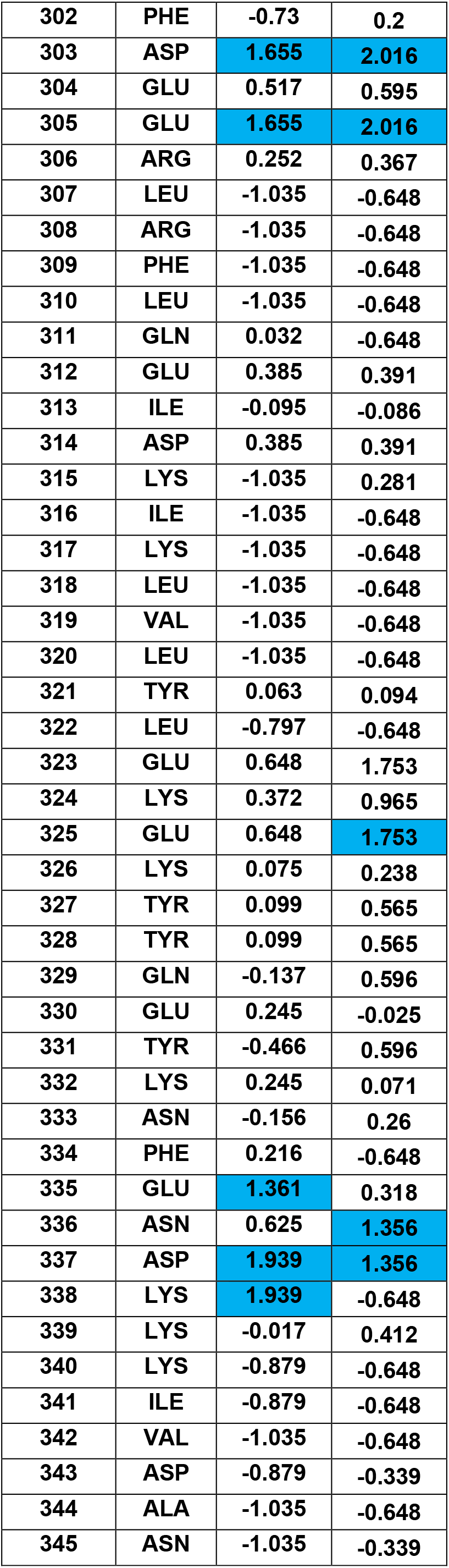
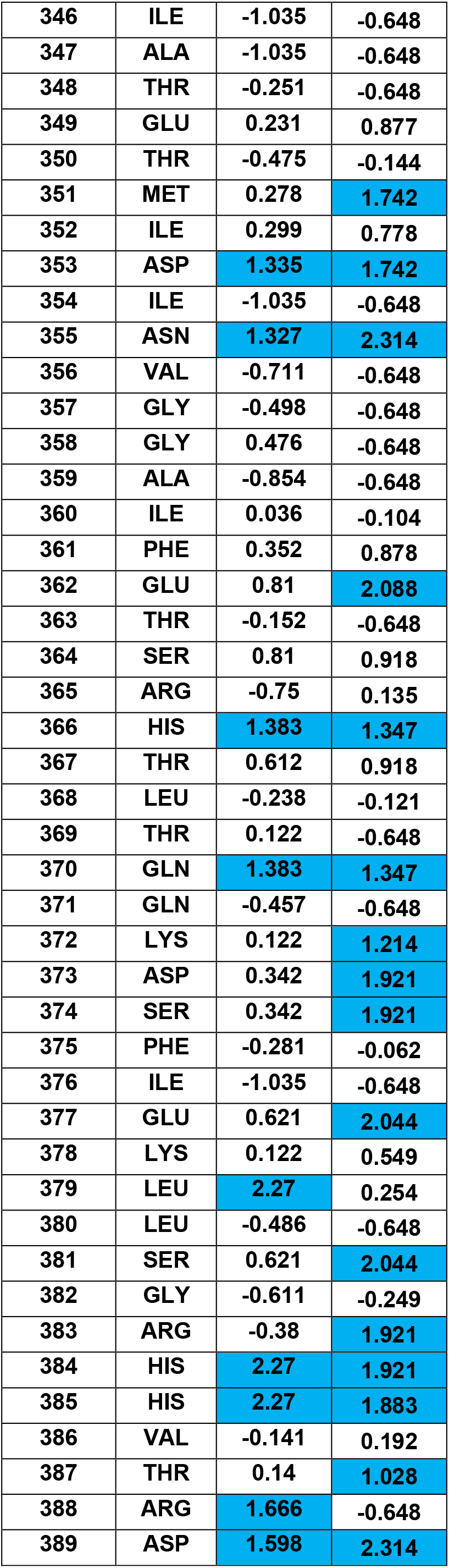
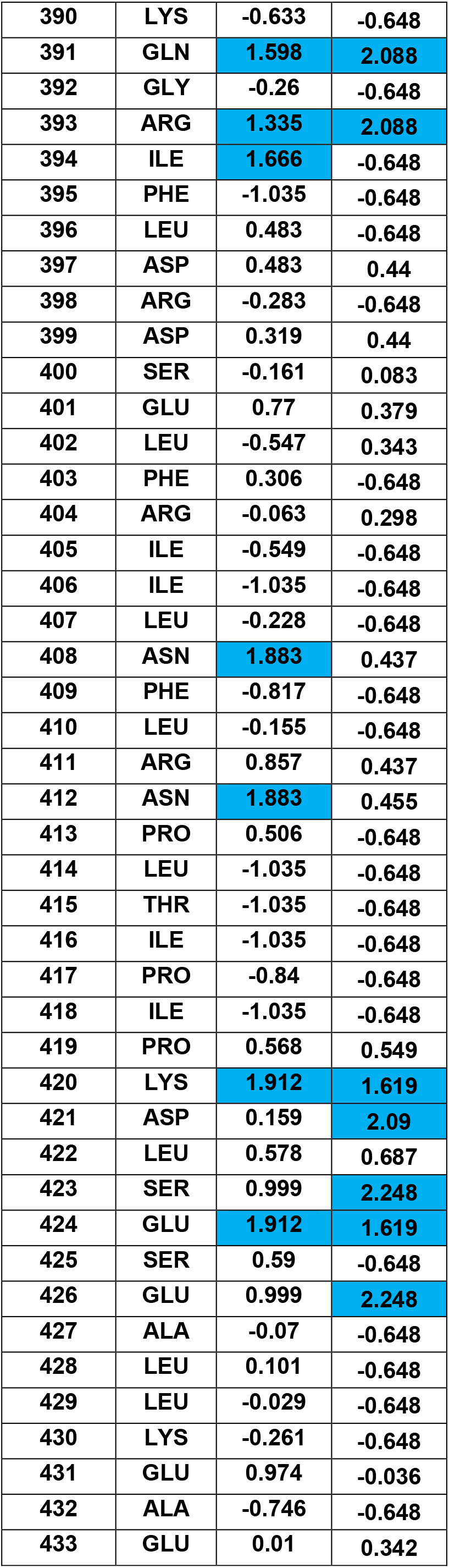
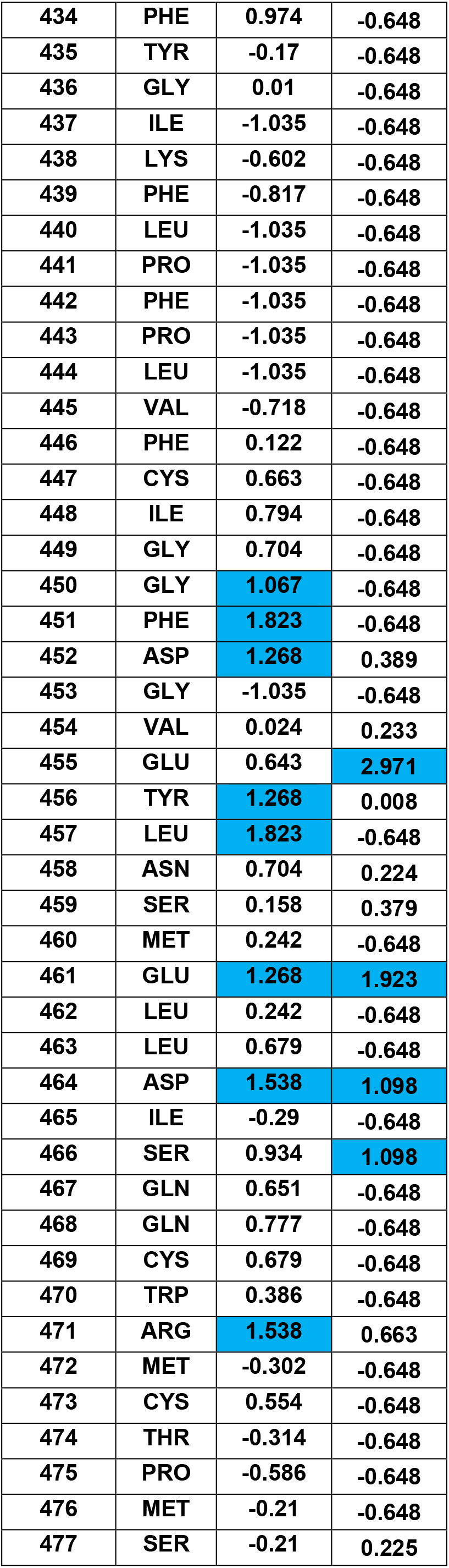
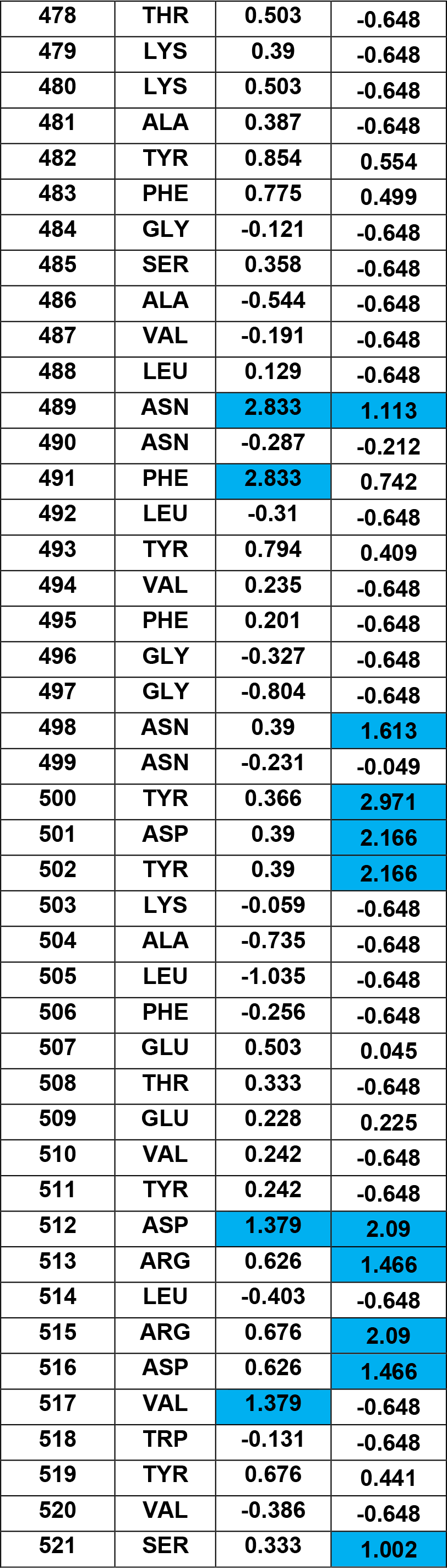
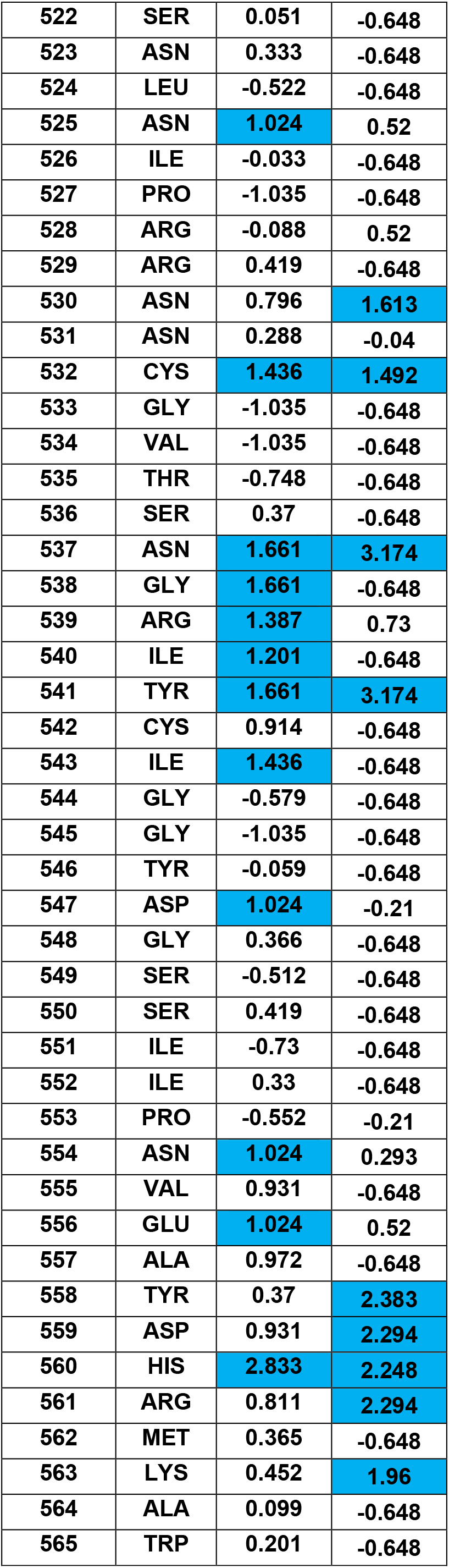
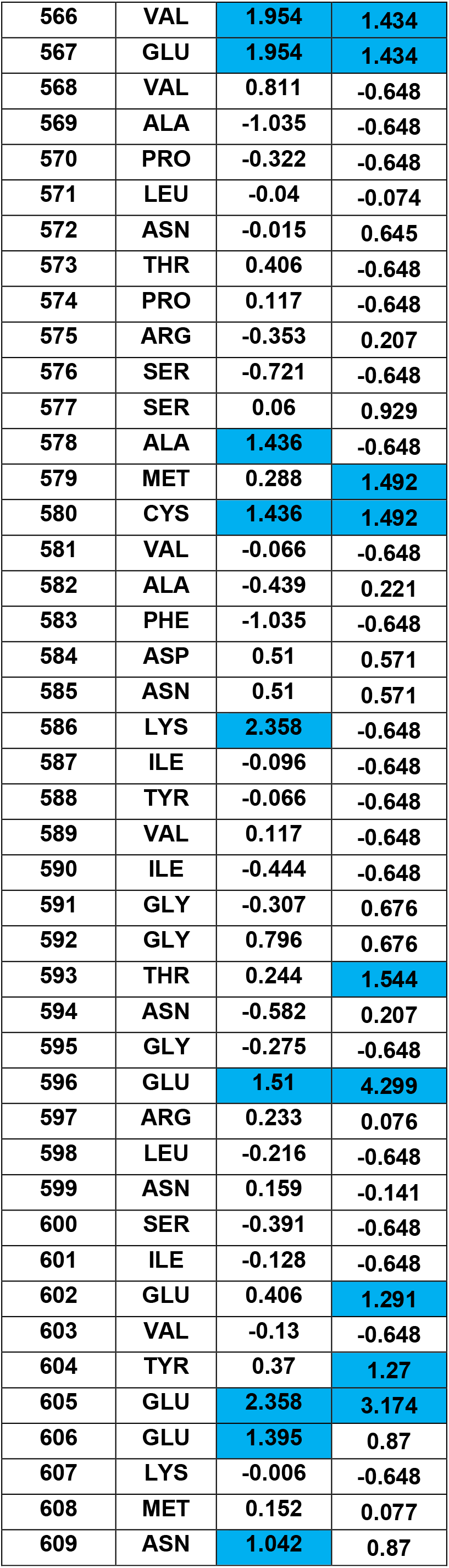
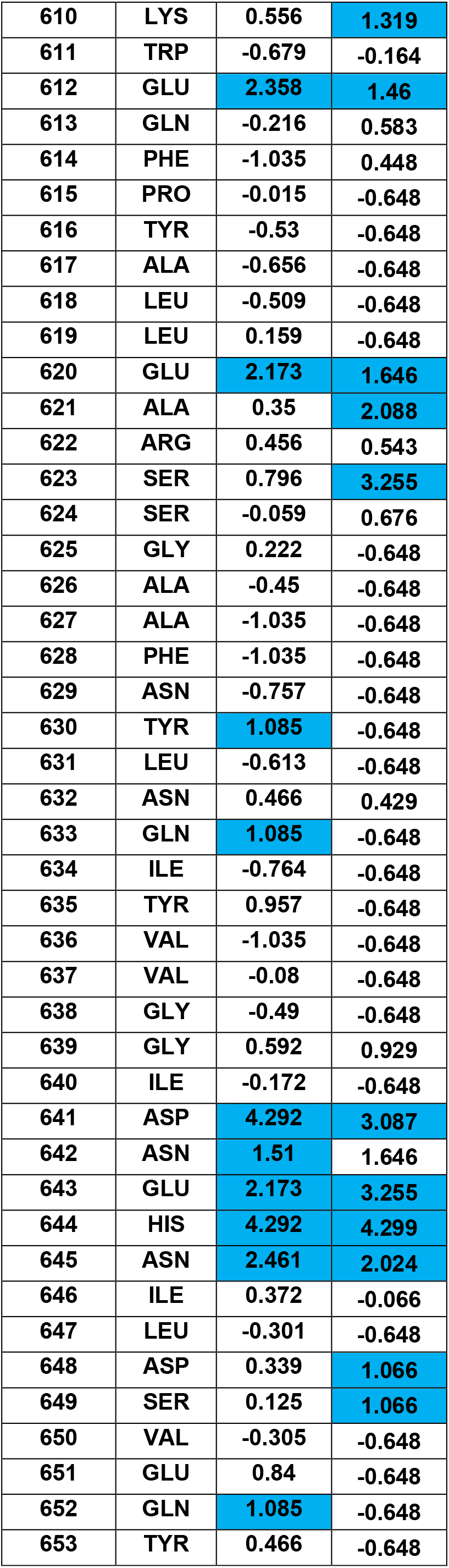
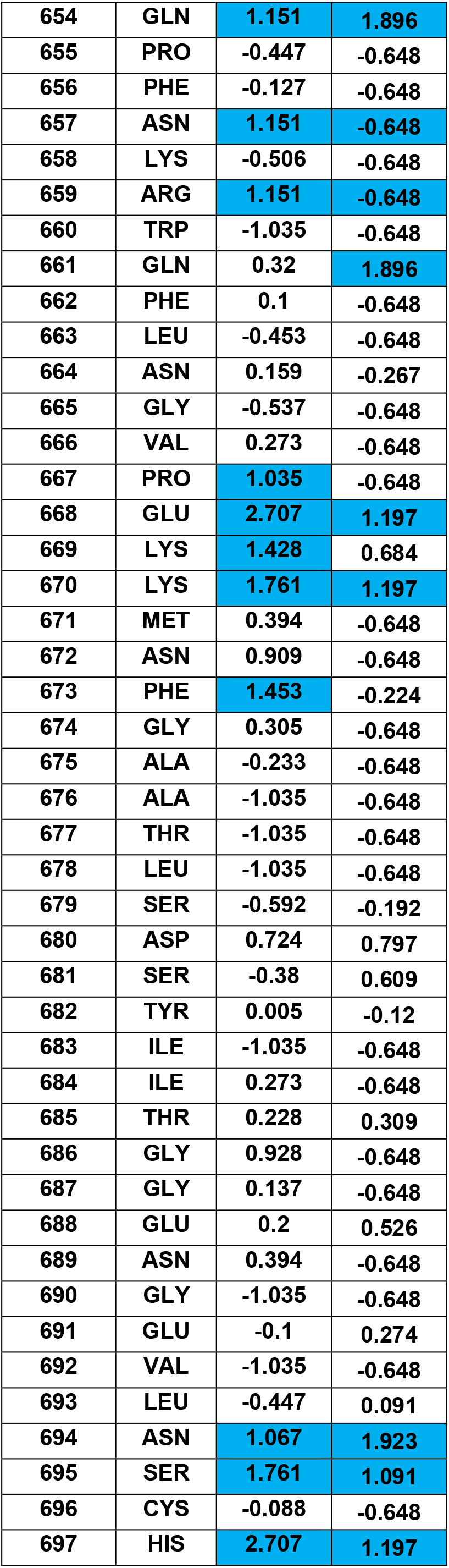
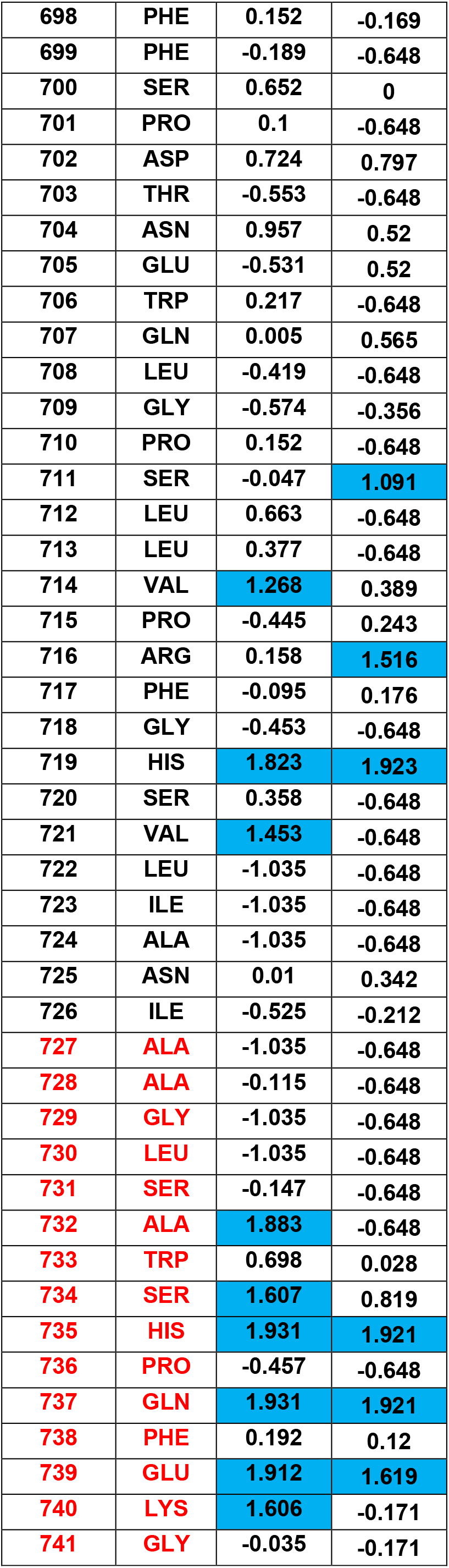
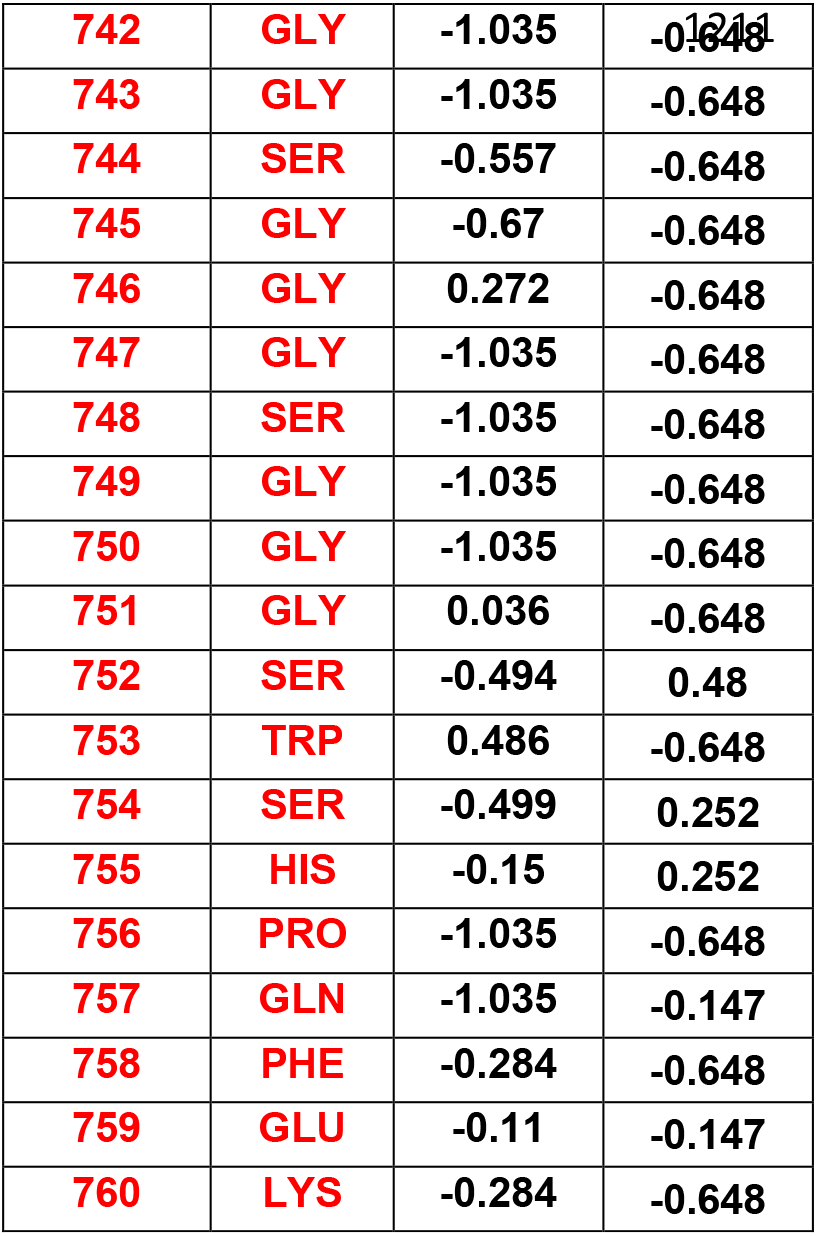
Fe2+ Binding Prediction score of the full length PfKelch13 with strep tag at the C-terminus (as according to http://combio.life.nctu.edu.tw/MIB2/structure). Amino acids from the C-terminus strep tag are in red text. Amino acids for the TrK13-WT protein are in bold. Predicted Fe2+ and Fe3+ binding residues with scores >1 are highlighted in blue.

**Supplementary Table S2:** Amino acid residues in PfKelch13 predicted to interact with heme. Amino acid residues in red and yellow text represent those which have either been validated or are candidate markers of ART-R, respectively102.

| WT and A578S | ID Pose | Affinity (kcal/mol) | Amino acids involved in the interactions |
| --- | --- | --- | --- |
|  | 20 | -7.9 | F451, G453, Y456, Y482, N498, R529, N530, S576, S577, R597, S623, S624, I640, I646, M671, N672, E688, G690, F717 |
|  | 14 | -7.5 | L422, R515, W518, Y519, V520, S521, S522, N523, Y558, K563, A564 |
|  | 9 | -7.1 | K586, Y588, V603, E605, E612, Q613, F614, P615, Y616, Y653, P655, K658 |
|  | 11 | -6.9 | E433, K438, V487, L488, N489, N490, T535, N537, C580, V581, A582, F583, D584, N585, N629, T677, I723, A724, N725 |
|  | 6 | -6.7 | Y502, K503, A504, F506, N523, L524, N525, I526, D547, G548, S549 |
|  | 12 | -6.4 | N572, T573, R575, N594, E596, R597, L598, N599, S600, I601, E602, Q613, F614, P615, Y616, A617, L618, N642 |
|  | 17 | -6 | L619, Q652, Q654, N657, R659, W660, Q661, F662, L663, N664, G665, N704 |
| **C580Y** | **ID Pose** | **Affinity (kcal/mol)** | **Amino acids involved in the interactions** |
|  | 18 | -8 | V534, T535, S536, **N537**, V581, A582, F583, D584, N585, E606, F628, N629, Y630, L631, N632, T677, L678, S679, D680, S681, Y682, P701 |
|  | 12 | -7.8 | F583, D584, K586, Y588, V603, Y604, E605, M608, K610, E612, Q613, F614, P615, Y616, N632, I634, Y653, P655, K658 |
|  | 5 | -7 | D452, G453, V454, Y456, Y482, N498, Y500, R529, N530, Y546, S576, S577, S624, I640, M671, N672, E688, G690, E691, F717 |
|  | 3 | -7 | V445, I448, **N458**, P475, **M476**, S477, T478, K479, N499, D501, Y502, L505, F506, E507, V520 |
|  | 6 | -6.9 | F491, **Y493**, V510, Y511, D512, **R515**, V517, Y519, V520, S521, S522, H560, K563 |
|  | 7 | -6.4 | N572, T573, R575, N594, E596, R597, L598, N599, S600, I601, E602, Q613, F614, P615, Y616, A617, A621 |
|  | 8 | -5.7 | **P553**, N554, E567, **V568**, A569, P570, L571, N572, N609, K610, W611 |
| **R539T** | **ID Pose** | **Affinity (kcal/mol)** | **Amino acids involved in the interactions** |
|  | 11 | -8 | V534, T535, S536, **N537**, V581, A582, F583, D584, N585, E606, F628, N629, Y630, L631, N632, T677, L678, S679, D680, S681, Y682, P701 |
|  | 8 | -7.7 | F583, D584, K586, Y588, V603, E605, M608, K610, E612, Q613, F614, P615, Y616, N632, I634, Y653, P655, K658 |
|  | 4 | -7.1 | I448, **Y482**, R529, N530, Y546, I551, S576, S577, T593, N594, G595, R597, S623, S624, I640, H644, M671, N672 |
|  | 6 | -7 | F491, **Y493**, V510, Y511, D512, **R515**, V517, Y519, V520, S521, S522, Y558, H560, K563 |
|  | 9 | -6.4 | S477, T478, K479, N499, D501, Y502, L505, F506, E507, V520 |
|  | 14 | -6.2 | N572, T573, R575, N594, E596, R597, L598, N599, S600, I601, E602, Q613, F614, P615, A617, A621 |
|  | 3 | -5.8 | N537, **T539**, Y541, A557, Y558, V566, E567, **V568**, Y604, E606, K607, M608, N609 |

## Supplementary Figure legends

**Supplementary Figure S1. Comparison of the KRP domain sequences of PfKelch13 and TaTFP with localization of the TaTFP positions involved in iron binding.** Sequence alignment between the KRP domains of PfKelch13 (PfK13) and TaTFP. The amino acid residues involved in Fe2+ binding in TaTFP are highlighted in yellow along with putative homologs in PfKelch13.

**Supplementary Figure S2. Expression of recombinant TrK13 proteins. A-B.** SDS-PAGE (A) and corresponding western blots (B) showing the selective induction of recombinant TrK13-WT, TrK13-R539T, TrK13-C580Y and TrK13-A578S at 61 kDa (arrowheads) only in the induced (I) samples but not in the uninduced (UI) lanes. For western blotting, custom-made anti-PfKelch13 antibodies were used. Molecular weight standards (in kDa) are as indicated.

**Supplementary Figure S3. Solubility of recombinant TrK13-WT protein in the presence of excess FAS or FES.** SDS-PAGE showing no change in the solubility status of TrK13-WT protein (10 µM) in the presence of 100-fold excess FAS or FES compared to the control. The TrK13-WT protein (5 µM) was incubated in the absence or presence of 500 µM iron salts for 30 mins, followed by centrifugation at 13,000 rpm for 30 min at 4°C to remove any aggregations. The supernatant fractions were resolved by SDS-PAGE and visualized by Coomassie staining. Molecular weight standards (in kDa) are as indicated.

**Supplementary Figure S4. UV-visible spectrophotometric scans. A-C.** UV-visible spectrophotometric scans showing the increase in absorbance profiles in the recombinant TrK13-C580Y (A), TrK13-A578S (B) and TrK13-R539T (C) proteins in the presence of both FAS (red) and FES (black). The standard absorbance profiles of these mutant TrK13 proteins without any metal salt are shown as black dotted lines.

**Supplementary Figure S5. Effect of chaotropic agent on dissociation of TrK13-WT oligomer and binding to FAS-NTA beads. A.** SDS-PAGE showing the stability of TrK13-WT oligomers at >180 kDa (arrow) up to 2 M urea and their gradual dissociation into the ∼61 kDa monomers (arrowhead) at higher urea concentrations, followed by glutaraldehyde cross-linking, as described earlier47. **B.** Relative abundance of TrK13-WT in the pulldown eluate with FAS-NTA beads from samples treated as in A. An approximately 50% reduction in relative abundance of eluted TrK13-WT was observed under conditions where the TrK13-WT oligomeric protein of >180 kDa disassembled into monomeric units of ∼61 kDa. Molecular weight standards (in kDa) are as indicated.

**Supplementary Figure S6.** Coomassie-stained SDS-PAGE (left) and graphical quantification (from three independent replicates, right) for the binding of the four recombinant TrK13 proteins to hemin-agarose beads (bottom left) relative to the respective input (top left). Molecular weight standards (in kDa) for SDS-PAGE are shown on the left.

**Supplementary Figure S7. Amino acid sequence of full-length PfKelch13**. Amino acid sequence of the full-length PfKelch13-WT-strep showing the location of the tryptophan (W) residues (bold, large font). The sequence of the recombinant TrK13-WT protein is underlined. The spacer and one-strep tag are shown in blue text.

**Supplementary Figure S8. Detection of heme-AS AAS by HPLC. A.** HPLC elution profiles showing the heme (tall peak at 8-10 mL) and AAS peaks (short peaks eluting around 15 ml) in reaction mixtures containing heme and AS. The AAS peaks are framed by dotted lines and magnified in **B.**

**Supplementary Figure S9. UV-visible spectrophotometric scans of heme, heme + ART, TrK13-WT + heme and TrK13-WT + ART. A.** Spectral scan showing no obvious change in the profiles of heme (2.5 µM) during the different incubation times. The characteristic Soret peak of heme at ∼400 nm is indicated by dotted lines. **B.** Spectral scans showing the time-dependent hypochromic and bathochromic/red-shift in the λmax from 400 nm to 412 nm in the absorbance of heme (2.5 µM) in the presence of ART (50 µM), likely due to heme-ART AAS formation. **C.** Spectrophotometric scans showing consistent and stable hypochromic and hypsochromic/blue-shifts in heme absorbance when bound to TrK13-WT protein. The spectrum of the TrK13-WT protein itself is shown as red dotted lines. The absorbance spectrum of free heme from panel B is also shown as a reference with black dotted lines. **D.** Spectral scan showing no visible time-dependent changes in TrK13-WT protein (10 µM) when incubated with 200 µM ART.

**Supplementary Figure S10. UV-visible spectrophotometric scans showing the absence of AAS formation by the TrK13-WT-bound heme analog Zn-PPIX.** Spectral scans showing the hyperchromic shift (red arrow) in the absorbance at 570 nm of the heme-TrK13-WT bound samples in the presence of ART. Blue dotted and solid lines represent scans before and after the addition of ART. Although the TrK13-WT protein binds the heme analog Zn-PPIX (green solid line), no apparent shift in absorbance is observed after the addition of ART (green dotted line), indicating no AAS formation. The spectrum of the TrK13-WT protein itself is shown as a dotted black line.

**Supplementary Figure S11. Expression and purification of recombinant *P. falciparum* Histidine-rich protein 2 (PfHRP2).** SDS-PAGE (left) and corresponding western blots (right) showing the selective induction of recombinant PfHRP2 at ∼68 kDa (arrowheads) in the induced (I) and Ni-NTA purified (P) lanes. Weak leaky expression is seen in the uninduced (UI) lanes. The western blot was performed with custom-made anti-PfHRP2 antibodies. The aberrant mobility of the recombinant PfHRP2 from its predicted molecular weight of 33.1 kDa is due to the presence of multiple histidine residues in the protein. Molecular weight standards (in kDa) are as indicated.

**Supplementary Figure S12. Pulldown of the native PfKelch13 protein from *P. falciparum* total protein extract with hemin-agarose beads.** 3D7 parasites were isolated by saponin treatment from asynchronized culture and proteins were solubilized with RIPA buffer. Protein extracts were incubated with hemin-agarose, or glutathione (GSH)-agarose as a control. After extensive washing, bound proteins were eluted with Laemmli buffer, separated by SDS-PAGE, and blotted against PfKelch13, and PfAldolase antibodies (black arrowheads). PfKelch13 antibodies were kindly provided by David Fidock36. The asterisk (*) below the 80-90-kDa band indicates lower molecular weight bands attributed to N-terminal degradation products of PfKelch1336. The parasite-derived PfKelch13 is shown to bind hemin-agarose beads (white arrowhead). Molecular weight standards (in kDa) are as indicated.

## Notes

### Competing Interest Statement

The authors have declared no competing interest.

