## Supplementary data for "Artemisinin-resistant *Plasmodium falciparum* Kelch13 mutant proteins display reduced heme-binding affinity and decreased artemisinin activation"

### Supplementary Figure S1

```
TaTFP_KRP      GGEDPPYESIDNDLYVFDFNTHTWSIAPANGDVPKTRVLGTRMVAVGTKLYVFGGRN---  95
PfK13_KRP      GGFDGV--EYLNSMELLDISQQCWRMCTPMST--KKAYFGS--AVLNNFLYVFGGNNYDY  502
               ** *    .  *.: ::*:. : * :.   .   *.  :*:  ..:.. ******.*

TaTFP_KRP      --------------------------------------------KQLEFEDFYSYDTVK  110
PfK13_KRP      KALFETEVYDRLRDVWYVSSNLNIPRRNNCGVTSNGRIYCIGGYDGSSIIPNVEAYDHRM  562
                                                             .  : :. :**

TaTFP_KRP      EEWKFLTKLDEKGGPEARTFHSMTSDENHVYVFGGVSKGGLNATPFRFRTIEAYNIAEGK  170
PfK13_KRP      KAWVEVAPLNTPR-----SSAMCVAFDNKIYVIGGTNGE-------RLNSIEVYEEKMNK  610
               : *  :: *:        :    .: :*::**:**..         *:.:**.*:   .*

TaTFP_KRP      WAQLPDPGEDFEKRGMAGFLVVQGKLWVFYGFATANDPKIPTLYGSQDYESNRVHCYDPA  230
PfK13_KRP      WEQFPYALLEARSSGAA--FNYLNQIYVVGGIDN------------EHNILDSVEQYQPF  656
               * *:*    : .. * *  :   .:::*. *: .            :.   : *. *:*

TaTFP_KRP      TQKWTEVETTGFEKPSRRSCFAHAAVGKYIIIFGGEIERDPEAHQGPGTLSREGFALDTE  290
PfK13_KRP      NKRWQFLNGV----PEKKMNFGAATLSDSYIITGGENGEVLNS----------CHFF---  699
               .::*  :: .    *.::  *. *::..  ** ***  .  ::           . :

TaTFP_KRP      TLVWERYEGGPIKPSNRGWVASTTTTINGKKGLLVHGGKLMTNERTDEMYFFAVNSST    348
PfK13_KRP      ------------SPDTNEWQLGPSLLVPR-FGHS----VLIANI--------------    726
                           .*... *  . :  :    *       *::*
```

**Supplementary Figure S2**

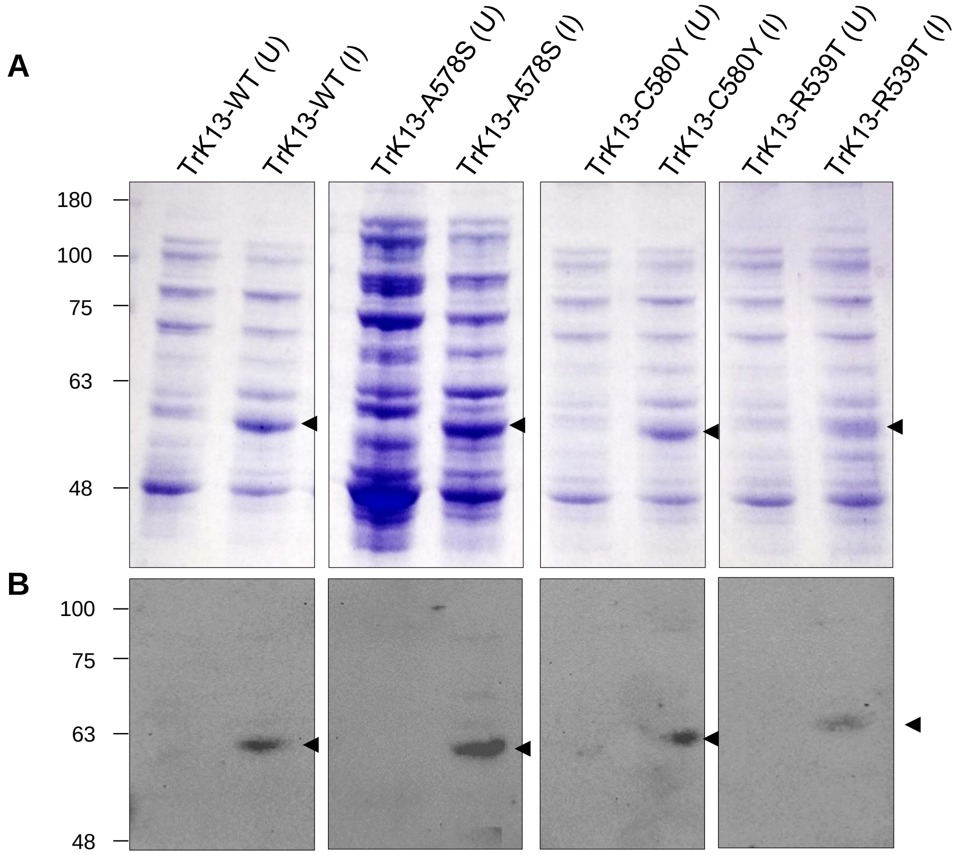

#### Supplementary Figure S3

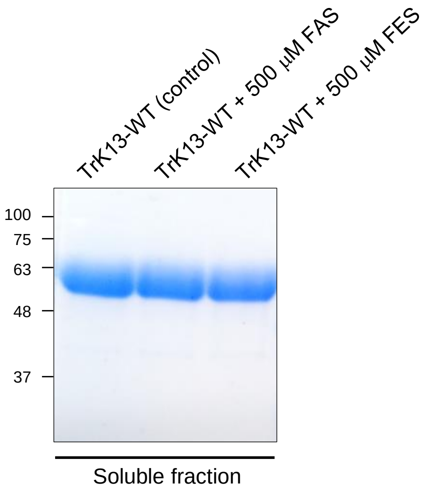

**Supplementary Figure S4**

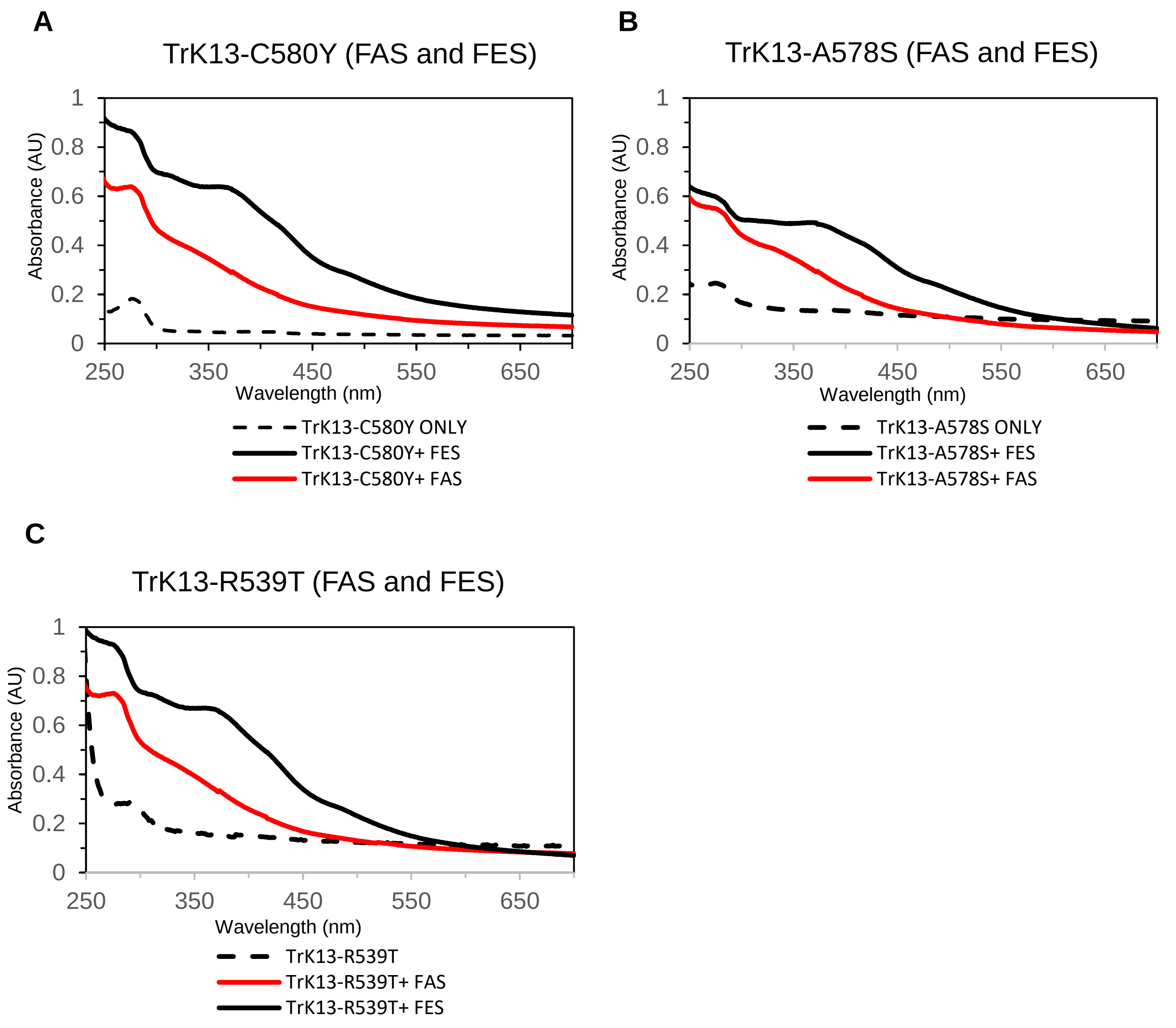

#### Supplementary Figure S5

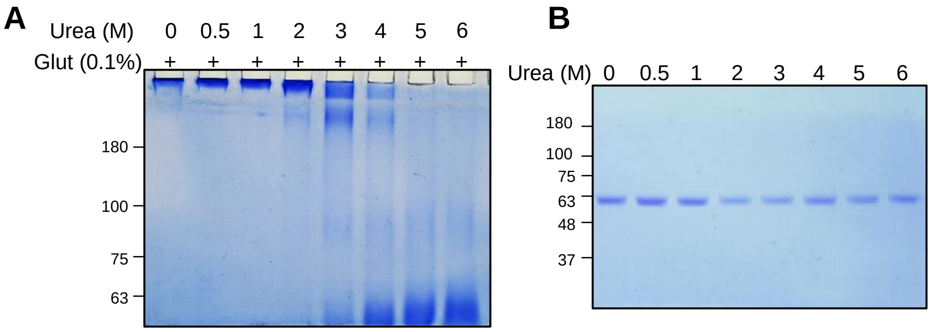

#### Supplementary Figure S6

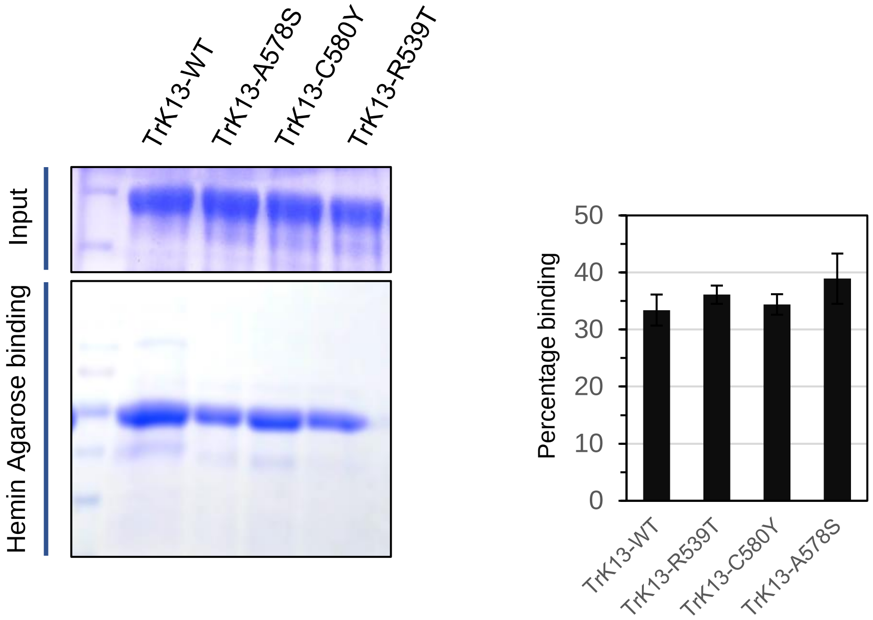

**Supplementary Figure S7**

MEGEKVKTKANSISNFSMTYDRESGGNSNSDDKSGSSSENDSNSFMNLTSDKNEKTENNSFLLN

NSSYGNVKDSLLESIDMSVLDSNFDSKKDFLPSNLSRTFNNMSKDNIGNKYLNKLLNKKKDTIT

NENNNINHNNNNNNLTANNITNNLINNNMNSPSIMNTNKKENFLDAANLINDDSGLNNLKKFST

VNNVNDTYEKKIIETELSDASDFENMVGDLRITFIN**W**LKKTQMNFIREKDKLFKDKKELEMER

VRLYKELENRKNIEEQKLHDERKKLDIDISNGYKQIKKEKEEHRKRFDEERLRFLQEIDKIKLV

LYLEKEKYYQEYKNFENDKKKIVDANIATETMIDINVGGAIFETSRHTLTQQKDSFIEKLLSGR

HHVTRDKQGRIFLDRDSELFRIILNFLRNPLTIPIPKDLSESEALLKEAEFYGIKFLPFPLVFC

IGGFDGVEYLNSMELLDISQQC**W**RMCTPMSTKKAYFGSAVLNNFLYVFGGNNYDYKALFETEV

YDRLRDV**W**YVSSNLNIPRRNNCGVTSNGRIYCIGGYDGSSIIPNVEAYDHRMKA**W**VEVAPLNT

PRSSAMCVAFDNKIYVIGGTNGERLNSIEVYEEKMNK**W**EQFPYALLEARSSGAAFNYLNQIYV

VGGIDNEHNILDSVEQYQPFNKR**W**QFLNGVPEKKMNFGAATLSDSYIITGGENGEVLNSCHFF

SPDTNE**W**QLGPSLLVPRFGHSVLIANIAAGLSA**W**SHPQFEKGGGSGGGSGGGS**W**SHPQFEK

#### Supplementary Figure S8

**A**

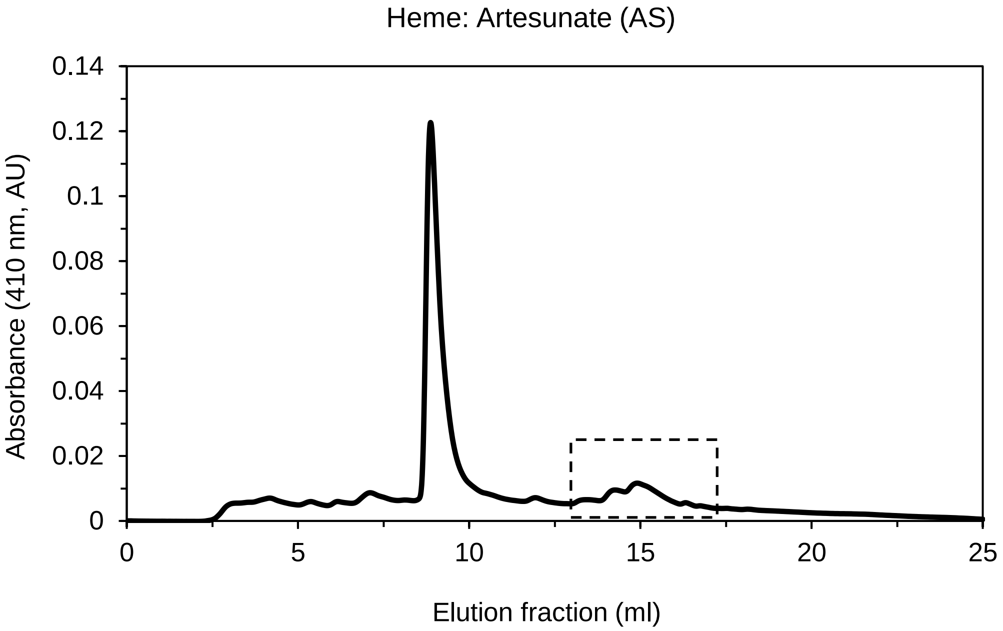


**B**

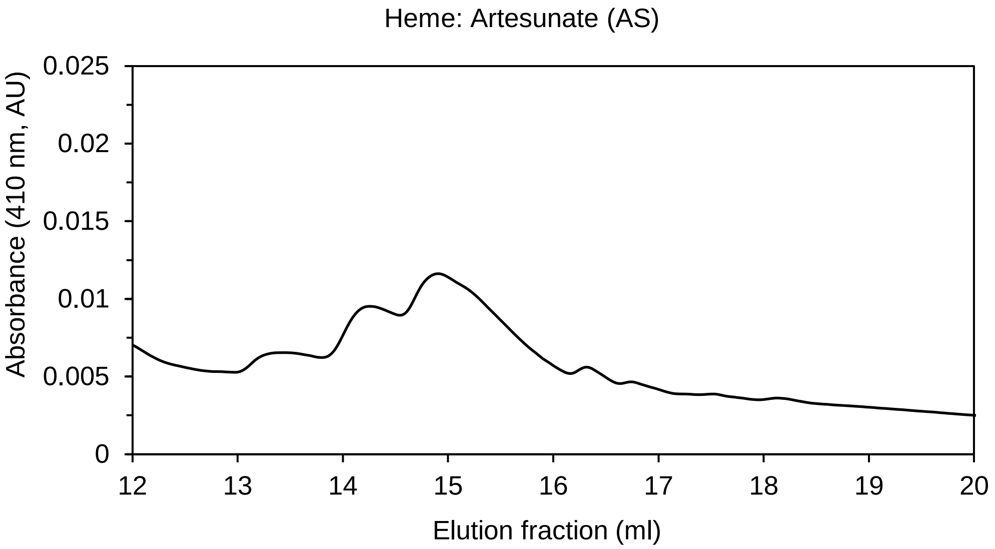

#### Supplementary Figure S9

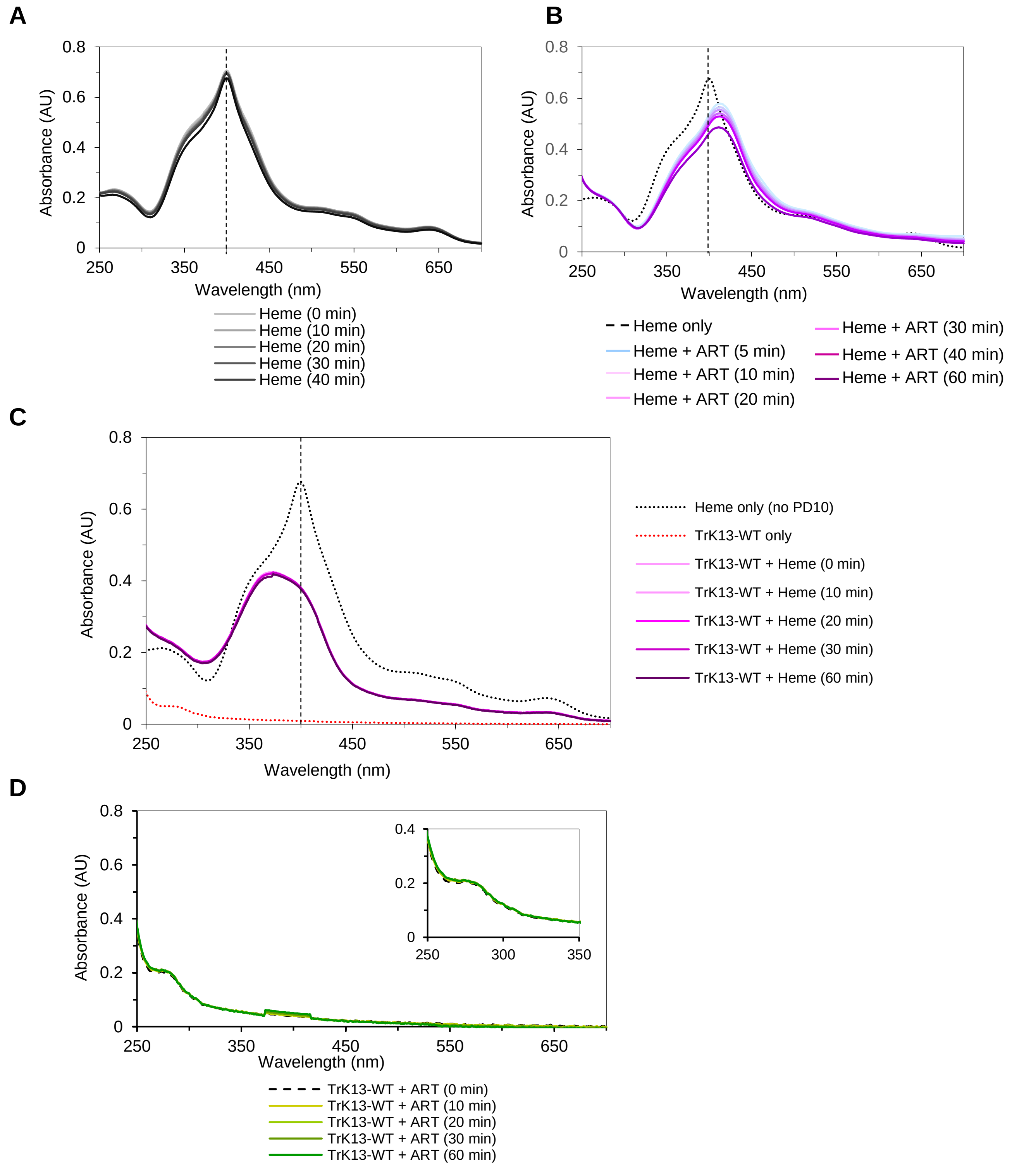

### Supplementary Figure S10

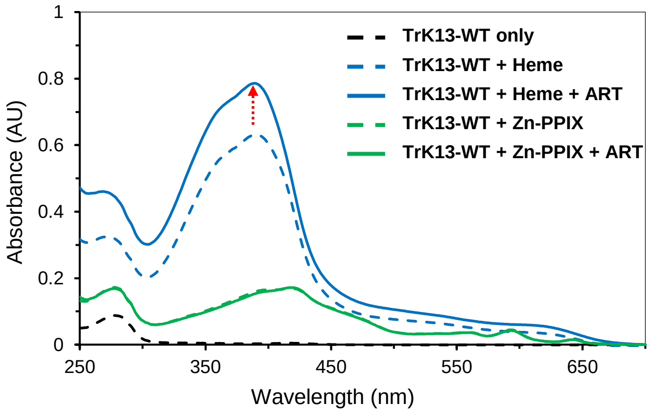

#### Supplementary Figure S11

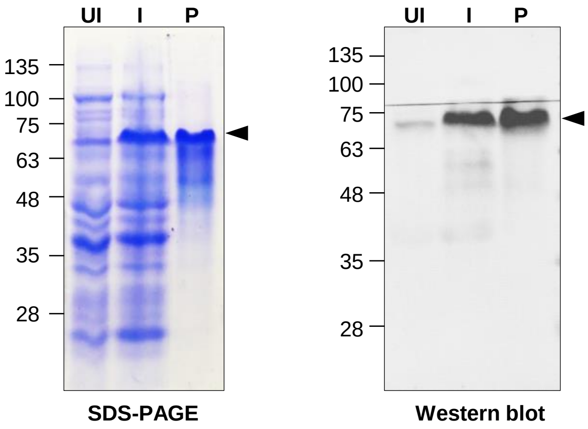

### Supplementary Figure S12

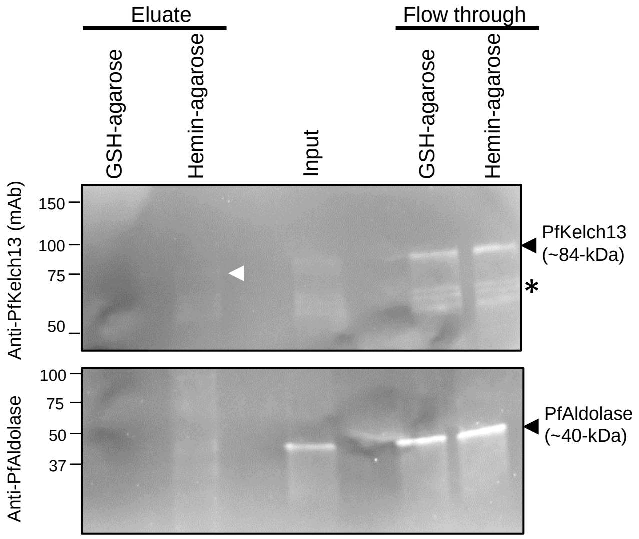
